# The ALOG domain defines a new family of plant-specific Transcription Factors acting during Arabidopsis flower development

**DOI:** 10.1101/2023.06.21.545689

**Authors:** Philippe Rieu, Veronica Beretta, Francesca Caselli, Emmanuel Thévénon, Jérémy Lucas, Mahmoud Rizk, Emanuela Franchini, Elisabetta Caporali, Max Nanao, Martin Kater, Renaud Dumas, Chloe Zubieta, François Parcy, Veronica Gregis

**Author notes:** These authors contributed equally (Philippe Rieu, Veronica Beretta). Structural Plant Biology Laboratory, Department of Botany and Plant Biology, University of Geneva, 1211, Geneva, Switzerland (Philippe Rieu). François Parcy and Veronica Gregis **Email:** and. **Author Contributions:** FP, PR, VG and MK developed the research project. PR performed the biochemical experiments helped by ET and RD. JL performed the bioinformatics analyses. PR and CZ performed the crystallization tests and CZ, MN and MZ acquired and analyzed the data. VG and EM generated the Arabidopsis *lsh* multiple mutants, VMB and FC performed the phenotypical characterization of *lsh* mutants, Y2H and confocal analyses, EC carried out SEM analyses. PR, VMB and FC assembled the figures. PR, FP, VG and VMB wrote the paper. All authors have read and agreed to the published version of the manuscript. **Competing Interest Statement:** There are no competing interests.

## Abstract

The ALOGs (Arabidopsis *LIGHT-DEPENDENT SHORT HYPOCOTYLS 1* and Oryza *G1*) are Transcription Factors (TFs) from an evolutionarily conserved plant-specific family shown to play critical roles in meristem identity, inflorescence architecture and organ boundaries in diverse species from mosses to higher flowering plants. However, the DNA binding-specificity and molecular determinants of protein-DNA interactions of this family were uncharacterized. Using *in vitro* genome-wide studies, we identified the conserved DNA motif bound by ALOG proteins from the liverwort *Marchantia polymorpha* and the flowering plants Arabidopsis, tomato and rice. In order to determine the amino acids responsible for DNA-binding specificity, we solved the 2.1Å structure of the ALOG DNA binding domain in complex with its cognate DNA. The ALOG DBD is an all-alpha helical domain with a structural zinc ribbon insertion and an N-terminal disordered NLS. The NLS sequence forms an integral part of the DNA binding domain and contributes to direct base read-out. To define the function of a group of redundant ALOG proteins in the model plant Arabidopsis thaliana, we generated a series of *alog* mutants and uncovered their participation in a gene regulatory network involving the other floral regulators LEAFY, BLADE-ON-PETIOLE and PUCHI, all active in defining boundary regions between reproductive meristems and repressing bracts development. Taken together, this work provides the biochemical and structural basis for DNA-binding specificity of an evolutionarily conserved TF family and reveals its role as a key player in defining organ boundaries in Arabidopsis.

**Significance Statement:** Transcription Factors (TFs) are key proteins that bind specific regions in the genome and regulate the expression of associated genes. Not all organisms possess the same set of TFs and some, like the ALOGs, are specific to the plant kingdom. These TFs have been shown to play important roles from mosses to flowering plants. However, it was not known what DNA motif they recognize and how they bind DNA. Here we identify this motif, we show it is widely conserved in evolution and we understand how this new type of DNA binding domain works at the structural level. In addition, we also show that several *ALOG* genes from Arabidopsis share a redundant function within the genetic network underlying correct floral meristem development.

## Introduction

The control of gene expression by Transcription Factors (TFs) is of key importance for all living organisms. Some TF types are widely shared among diverse organisms whereas others have emerged in specific groups. Plants, in particular, possess a diversity of TFs that are absent in other kingdoms. Many of these plant-specific TF families were born around the time of emergence from the water (in charophytes algae or mosses) and have later expanded and been co-opted in various processes along plant evolution (1). Most of these plant-specific TFs have been characterized at the biochemical and structural levels. Still, a few proteins identified as important regulators of plant physiology or development are suspected to act as TFs, but the characterization of their DNA binding specificity or the structural basis for DNA recognition is still missing. This is the case for the *ALOG* (*Arabidopsis LIGHT-DEPENDENT SHORT HYPOCOTYLS 1 (LSH1) and Oryza G1*) gene family. These genes, first described in Arabidopsis (2) are present in land plants as well as in some algae (3, 4), usually as gene families. For example, the *Arabidopsis thaliana* genome contains 10 *LSH* genes (2).

*ALOG* genes have shown to play crucial developmental roles in diverse plant species. In the liverwort *Marchantia polymorpha, LATERAL ORGAN SUPPRESSOR 1* (*LOS1*) and *LOS2* are implicated in meristem maintenance and lateral organ development (4, 5). In tomato, the *ALOG* gene *TERMINATING FLOWER* (*TMF*) controls flowering by preventing the precocious expression of *ANANTHA (*the tomato ortholog of the Arabidopsis the *UNUSUAL FLORAL ORGAN* (*UFO*) gene) (*AN*) (6). In rice, the *ALOG* genes *LONG STERILE LEMMA1* (*G1*) (7), *TRIANGULAR HULL 1* (*TH1*) (8), *TAWAWA1* (*TAW1*) (9) and *G1-LIKE 1* and *G1-LIKE 2* (*G1L1* and *G1L2*) (10) have shown to play diverse roles, such as controlling sterile lemma, lemma, palea and panicle development. In pea, the *ALOG* gene *SYMMETRIC PETALS 1* (*SYP1*) is a regulator of floral organ internal asymmetry (11) and in *Medicago truncatula*, LSH1/LSH2 control nodule identity (12). In Arabidopsis, the role of *LSH* genes have been studied, in particular LSH8 (13), LSH9 (14) and LSH10 (15). Mutants have also been isolated for *LSH1, LSH3* and *LSH4* but none display any obvious phenotype (16, 17). However, constitutive expression of *LSH4* (and to a lesser extent *LSH3*) results in the inhibition of leaf growth, the production of extra floral organs, chimeric floral organs, or shoots within a flower (17). *LSH3* and *LSH4* are active in the boundary regions of shoot organs under the transcriptional control of the CUP-SHAPED COTYLEDON1&2 TFs (17, 18). Overall, these phenotypes suggest that Arabidopsis LSH3 and LSH4 might repress organ formation in boundary regions and control floral meristem identity acquisition.

Despite this increasing body of evidence that ALOG proteins play important roles in various plant species, little is known about the molecular and structural properties of these TFs. Several ALOG proteins were shown to directly interact with promoter regions and act as transcriptional repressors (15, 19, 20). A well-studied example is the repression of the tomato *AN* gene by the ALOG protein TMF, involving phase separation on the *AN* promoter (19, 21). At the molecular level, ALOG proteins share a conserved domain (named ALOG domain) (3), flanked by non-conserved disordered regions of variable lengths. The ALOG domain likely originates from the DBD of bacterial recombinases found in mobile elements (3), with a putative zinc ribbon inserted between predicted helices 2 and 3. Which structural elements (helices and/or zinc ribbon) participate in sequence-specific DNA binding was not known.

Here, we identify the ALOG-bound DNA motif using *in vitro* genomic binding assays and show that this motif is recognised by ALOG proteins from moss to flowering plants, demonstrating a high evolutionary conservation of the domain and recognition sequence. This motif allowed us to solve the structure of the ALOG DNA-binding domain of LSH3 from *Arabidopsis thaliana* in complex with its cognate DNA and determine the structural properties of a new family of plant-specific TFs. We also deepen our understanding of the role of *LSH1, LSH3* and *LSH4* in the reproductive development of *Arabidopsis thaliana*. Our genetic and molecular analyses show that these three genes have overlapping functions in defining specific identities and boundaries within Arabidopsis flower meristems. Taken together, these studies demonstrate that ALOG proteins act as TFs with a high evolutionary conservation of structure and DNA-binding specificity with roles in reproductive structures and developmental transitions.

## Results

### Determination of the ALOG DNA-binding specificity

We studied ALOG TFs from several plants (Fig. 1A), selecting those with available genetic or biochemical data (2, 4–6, 17). We included OsG1 as it contains two unique insertions, absent from other ALOG proteins (7). To establish their DNA binding specificity, we performed amplified DNA Affinity Purification sequencing (ampDAP-seq) (22) using Full Length (FL) *in vitro*-produced ALOG proteins (from Arabidopsis, tomato, rice and Marchantia) and Arabidopsis genomic DNA. AmpDAP-seq was performed in triplicates and yielded highly reproducible results (SI Appendix, Fig. S1). Motif search in ALOG-bound genomic regions robustly identified the same 7-bp YACTGTW (Y=T/C, W=A/T) motif for all the tested proteins (Fig. 1B and SI Appendix, Fig. S1), showing that ALOG DNA binding specificity is likely conserved throughout land plants. This motif displays several high-information positions, strongly suggesting specific contacts between ALOG residues and DNA bases at these locations. The Position Weight Matrix (PWM) corresponding to this motif reliably predicted ALOG DNA binding (Fig. 1C and SI Appendix, Fig. S1). The motif showed little symmetry, suggesting it is bound by an ALOG monomer.

**Figure 1.**
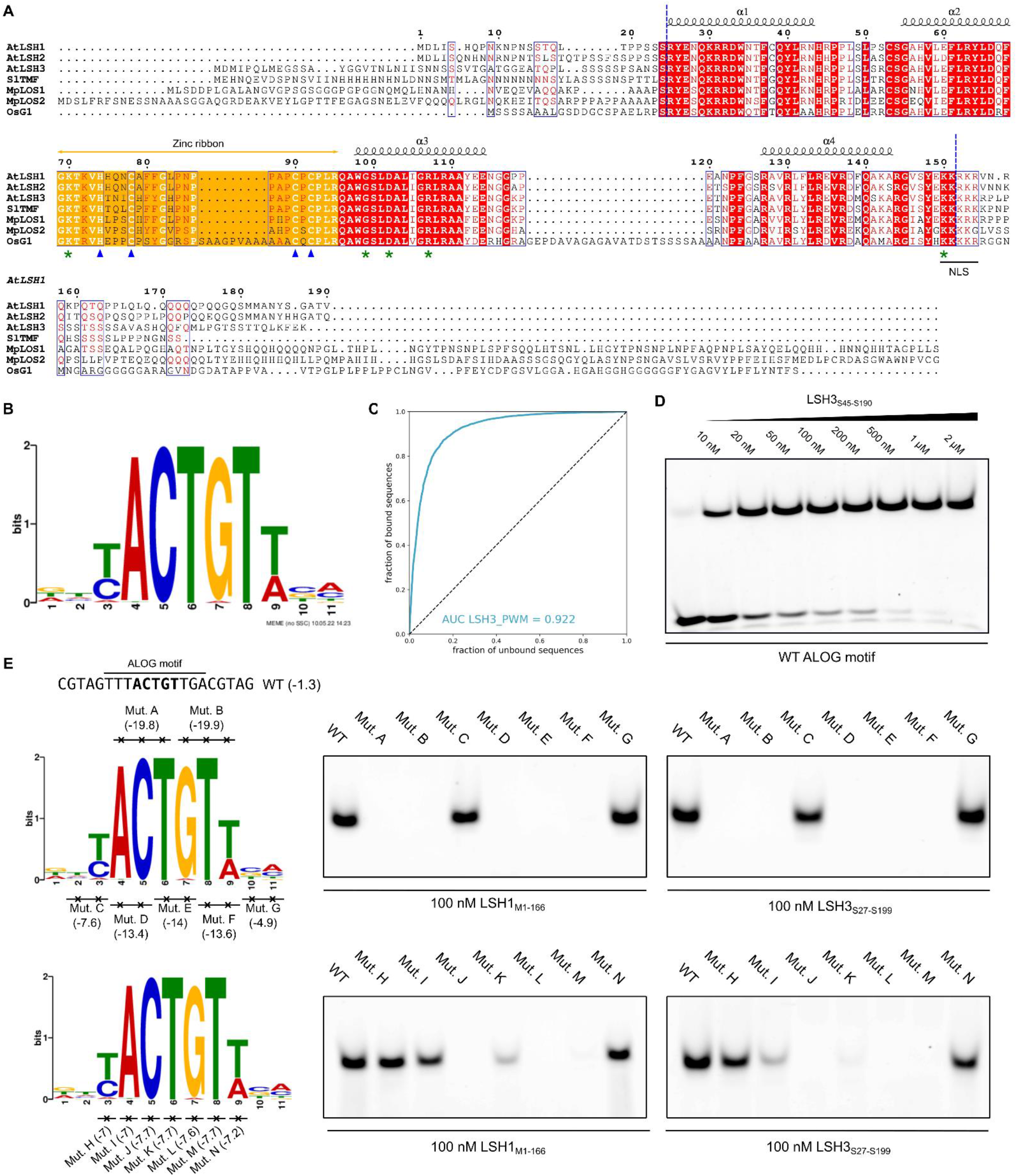
Determination of ALOG binding specificity by ampDAP-seq. (**A**) Alignment of studied ALOG proteins. Blue dotted lines indicate the limits of the ALOG domain. The zinc ribbon is highlighted in orange, with the key histidine and cysteine residues indicated by blue triangles. Green stars denote residues in direct contact with DNA bases (see Fig. 2). NLS = Nuclear Localization Signal. Numbers are relative to AtLSH1. At = *Arabidopsis thaliana*, Sl = *Solanum lycopersicum*, Mp = *Marchantia polymorpha*, Os = *Oryza Sativa*. (**B**) Logo obtained for LSH3 in ampDAP-seq, using the 600 peaks with the strongest signal. (**C**) Receiver operating characteristic (ROC) curve for LSH3 using all peaks except those used to build the logo. The value of the Area Under the Curve (AUC) is indicated. (**D**) EMSA with ALOG highest-score sequence DNA probe (WT ALOG motif) and LSH3_S45-S190_. Based on the analysis of 3 independent EMSAs, we found an apparent Kd of 28 nM for LSH3-DBD/DNA. (**E**) EMSA with LSH1_M1-L166_, LSH3_S27-S199_ and indicated DNA probes. The WT ALOG probe (top left) was mutated at positions indicated on the LSH3 logos (bottom left). Scores between brackets were obtained by scanning each DNA probe sequence with the LSH3 PWM (the best binding sites correspond to the less negative score values). EMSA with described DNA probes (right). Uncropped gels are provided in Dataset S3.

Next, we validated the ampDAP-seq motif using Electrophoretic Mobility Shift Assay (EMSA). We found that the *in vitro*-produced FL proteins and the isolated ALOG domains behaved similarly (SI Appendix, Fig. S2A), showing that the binding specificity is fully conferred by the ALOG domain. The ALOG DNA Binding Domain (DBD) was thus used in the rest of this work. EMSA using a DNA probe of optimal affinity according to LSH3 PWM showed a single shifted band with LSH1 or LSH3 DBDs (Fig. 1D and SI Appendix, Fig. S2B) with apparent Kds for LSH1 and LSH3 DBDs estimated below 50 nM, in the range of other affinities found for TF/DNA (23). Bases of this motif were systematically mutated and, overall, most mutations at the high-information positions of the motif strongly reduced binding of LSH1 and LSH3 DBDs (Fig. 1E). These experiments validated the YACTGTW motif as the ALOG binding site in land plants.

### Biochemical and structural characterisation of ALOG DBD

Based on the ampDAP-seq experiments, the ALOG proteins are monomeric unlike some previous reports of dimer formation by ALOGs (20, 24). We confirmed the oligomerization state of the FL ALOG proteins by co-immunoprecipitation (SI Appendix, Fig. S2C), Yeast-Two-Hybrid (Y2H: SI Appendix, Fig. S2D) and EMSA (SI Appendix, Fig. S2E), all of which demonstrated only monomeric species. As the DBD is sufficient for DNA-binding specificity, we structurally characterized the DBD of LSH3 in complex with the ampDAP-seq derived YACTGTW motif. Of note, the minimal DBD included the conserved Nuclear Localization Signal (NLS; Fig. 1A), which is required for DNA binding (SI Appendix, Fig. S3A and B). The LSH3 DBD was expressed in bacteria and purified to homogeneity by size exclusion chromatography and subjected to crystallization trials in the presence of its cognate DNA. Crystals obtained with LSH3 DBD (LSH3_S45-S190_) bound to the YACTGTW motif (TAGTT**TACTGTT**GACGT DNA molecule) yielded a structure of the complex at 2.1 Å resolution (Table 1). We found that the ALOG DBD is primarily alpha helical, consisting of 4 alpha helices (with helices 1, 3 and 4 contacting the DNA major groove) and a C-terminal extension (comprising the NLS) providing additional direct DNA contacts (Fig. 2A-D). As previously proposed (3), the overall fold bears a high degree of structural similarity (conservation of the core arrangement of alpha helices) to DBDs from LoxP recombinase (PDB 1NZB), tyrosine recombinase (PDB 5HXY) and the apoptosis regulator protein BCL-2 (PDB 6YLD) (25). However, unlike its closest structural homologs, the ALOG domain has a 24 amino acid loop insertion between helices 2 and 3. A Zn^2+^ ion, chelated by three cysteines and a histidine residue (103-HxxxC…119-CxC) donated from this loop, forms a non-canonical zinc ribbon-like arrangement similar to a Gag knuckle (Fig. 2F) (26).

**Table 1.** Data collection and refinement statistics.

|  | ALOG-DNA |
| --- | --- |
| **Data collection** |  |
| Space group | P4<sub>3</sub>22 |
| Cell dimensions |  |
| *a*, *b*, *c* (Å) | 79.80, 79.80, 127.01 |
| α, β, γ (°) | 90, 90, 90 |
| Resolution (Å) | 67.-2.14 (2.19-2.14)* |
| *R*<sub>sym</sub> or *R*<sub>merge</sub> (%) | 4.5 (223) |
| *I* / σ*I* | 21.7 (0.9) |
| Completeness (%) | 99.8 (99.1) |
| Redundancy | 7.0 (6.9) |
| CC(1/2) | 100 (32.5) |
| **Refinement** |  |
| Resolution (Å) | 50.-2.14 (2.19-2.14) |
| No. reflections | 43003 |
| *R*<sub>work</sub> / *R*<sub>free</sub> | 21.4/25.2 (40.1/42.85) |
| No. atoms | 1809 |
| Protein | 1402 |
| Water | 51 |
| DNA | 345 |
| Zn<sup>+2</sup> | 1 |
| *B*-factors |  |
| Protein | 67 |
| Water | 67 |
| DNA | 92 |
| Zn<sup>2+</sup> | 59 |
| R.m.s. deviations |  |
| Bond lengths (Å) | 0.003 |
| Bond angles (°) | 0.54 |

\* refers to the highest resolution shell

**Figure 2.**
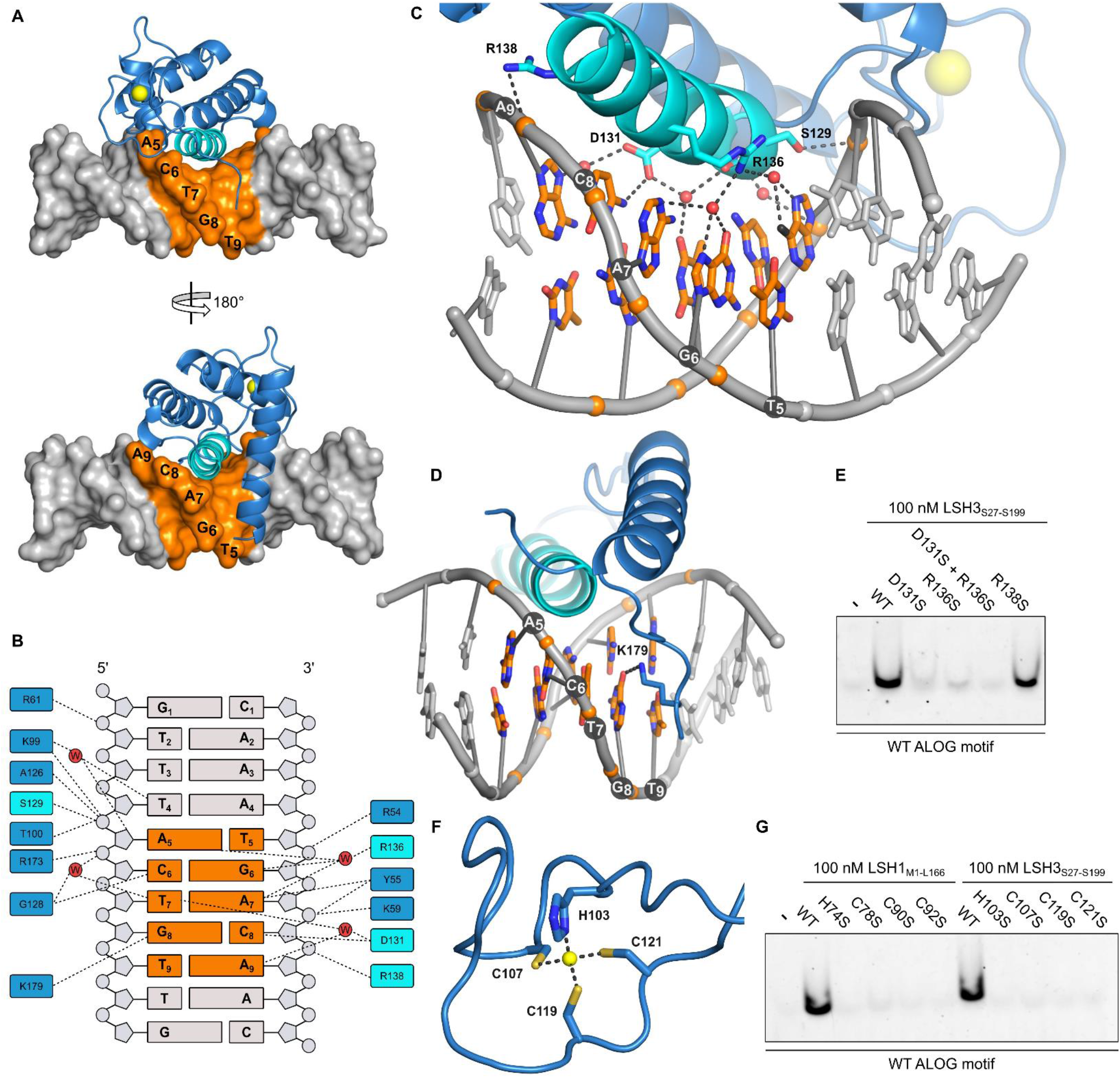
Structure of LSH3 DBD in complex with DNA. (**A**) ALOG-DBD/DNA complex. Throughout this figure, LSH3 DBD is shown in blue except helix 3 colored in cyan. The DNA duplex is depicted in gray with bases with the highest information content in orange. The Zn^2+^ ion is shown in yellow. A 180° rotation along the y-axis was applied to obtain the bottom picture. (**B**) Protein-DNA interactions. (**C**) Ribbon diagram of LSH3 DBD bound to its cognate DNA, with a focus on helix 3. Interactions are indicated by black dashed lines. For clarity, helix 1 was removed. (**D**) Close-up view of LSH3 C-terminal tail in contact with DNA. (**E**) EMSA with ALOG highest-score sequence DNA probe (WT ALOG motif) and indicated LSH3_S27-S190_ versions. (**F**) Close up view of the zinc ribbon. Side chains of the residues coordinating the zinc ion are represented. (**G**) EMSA with WT ALOG motif DNA probe and indicated LSH3_S27-S190_ and LSH1_M1-L166_ versions. Uncropped gels are provided in Dataset S3.

Point mutations in helix 3 (D131S and R136S), which make direct contacts with DNA bases (Fig. 2B and C) resulted in dramatically reduced DNA binding (Fig. 2E). Mutations of the residues from helices 1 and 4 that contact the phosphate backbone but are not involved in direct base read-out also somewhat reduced DNA binding (SI Appendix, Fig. S3C-E). Mutating the Zn^2+^-coordinating residues (Fig. 2G) or deleting LSH1 or TMF zinc ribbons to mimic recombinases also abolished binding to the ALOG motif (SI Appendix, Fig. S3F), confirming the importance of the Zn ribbon as a structural component of the protein (19). Based on structural comparisons to all other plant TF-DNA complexes, we propose that the ALOG domain defines a new family of plant TFs within the Zn coordinating DBD superclass and the HC3 class of Plant-TFClass (27, 28).

### *LSH1, LSH3* and *LSH4* functional characterization in *Arabidopsis thaliana* reproductive meristems

Next, we investigated the role of *LSH* genes in Arabidopsis. The expression of *LSH1-4* in inflorescence tissue and overexpression experiments suggested they could play a role in meristem boundaries determination and organogenesis (16–18). According to published expression data, *LSH1, LSH3* and *LSH4* have similar expression profiles in the IM, FMs and Stage-3 (ST3) flower tissues while the expression of *LSH2*, their closest homolog, is almost undetectable in those tissues (SI Appendix, Fig. S4). We further characterized the spatial expression profile of *LSH1, LSH3* and *LSH4* using *in situ* hybridization. Consistent with *LSH3* and *LSH4* published data (17, 29), all three genes were expressed in the boundary region between the IM and the FM. *LSH1* and *LSH3* were also expressed in the cryptic bract region while *LSH1* expression further extends in the IM (Fig. 3A-F). This latter observation suggests possible functional redundancy among the similarly expressed *LSH1, LSH3* and *LSH4* genes as the *lsh1, lsh3* or *lsh4* single mutants did not display any obvious phenotype (16–18).

**Figure 3.**
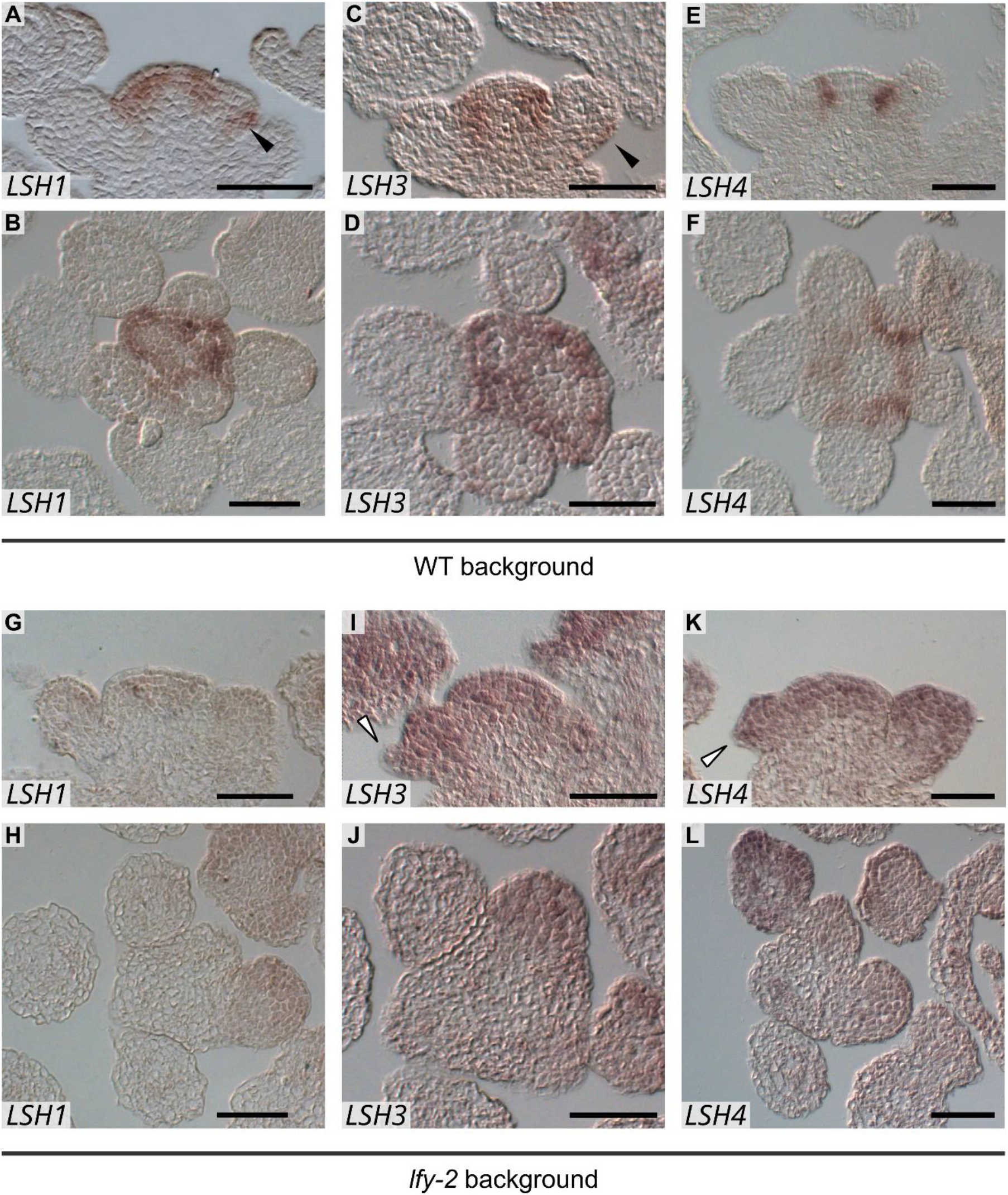
Expression of LSH RNAs in WT and *leafy-2* (*lfy-2*) mutant reproductive meristems. Expression profile of *LSH1* (A, B, G, H), *LSH3* (C, D, I, J) and *LSH4* (E, F, K, L) in reproductive tissues analyzed by *in situ* hybridization in WT (A-F) and *lfy-2* (G-L) backgrounds. A, C, E, G, I, K are longitudinal and B, D, F, H, J, L are transversal sections (scale bars = 50 μm). Black arrowheads in A and C indicate signal in the cryptic bract region, white arrowheads in I and K indicate bract primordia. See SI Appendix, Fig. S10 for negative controls.

To unravel their role, we generated novel alleles of *lsh1, lsh3* and *lsh4* mutants using the CRISPR-Cas9 gene editing system. For all three genes, we obtained mutations with a premature stop codon leading to truncated proteins lacking the ALOG domain (SI Appendix, Fig. S5). None of the single (*lsh1, lsh3* and *lsh4)* nor double mutants *(lsh1 lsh3, lsh1 lsh4, lsh3 lsh4)* display any visible aberrant phenotype. We thus analyzed the *lsh1 lsh3 lsh4* triple mutant and observed flowers subtended by a well-developed bract-like organ emerging from the pedicel. Such organs do normally not develop in wild type (WT) Arabidopsis plants (Fig. 4A and B). The triple mutant flowers did not show any other aberrations: all four types of floral organs had normal numbers, shapes and dimensions and the flowers were fully fertile. Scanning Electron Microscopy (SEM) analysis of *lsh1 lsh3 lsh4* bracts showed that these organs display sepal-like cell types and occasionally develop structures resembling stigmatic papillae (Fig. 4C-F).

**Figure 4.**
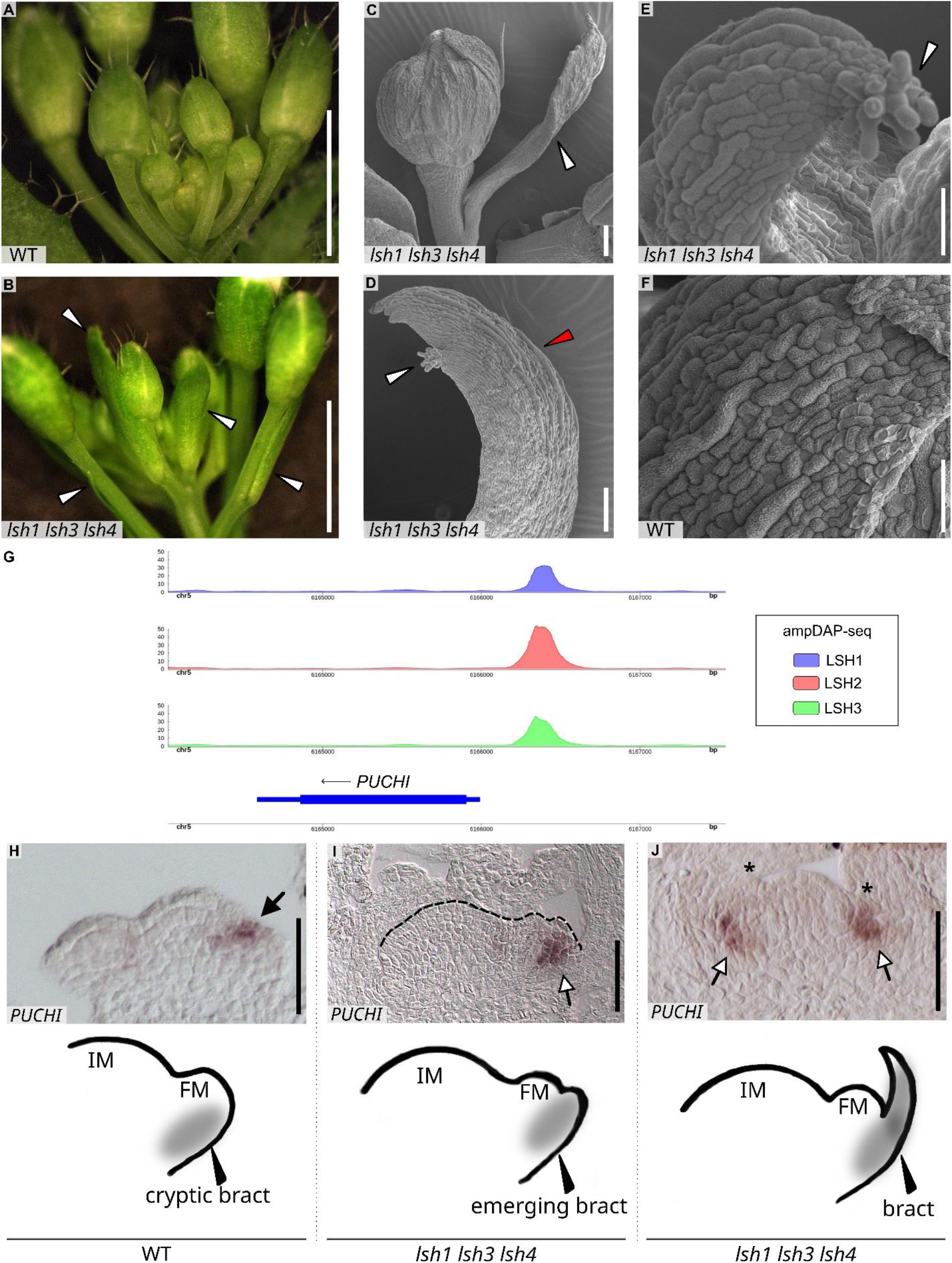
Analysis of the *lsh1 lsh3 lsh4* triple mutant. (**A, B**) Stereo microscope pictures of WT (A) and *lsh1 lsh3 lsh4* triple mutant (B) inflorescences. The white arrowheads in B indicate bracts (scale bars = 3 mm). (**C-F**) Scanning electron microscope (SEM) pictures of WT and *lsh1 lsh3 lsh4* plants. C, floral bud, the white arrowhead indicates the bract (scale bar = 100 µm). *lsh1 lsh3 lsh4* triple mutant bract (D and E). The white arrowhead indicates the stigmatic papillae. The red arrowhead indicates the sepal-like cells (scale bars: D = 100 µm; E = 50 µm). WT sepal (F, scale bar = 50 µm). (**G**) Genome browser view showing ampDAP-seq binding of LSH proteins to the *PUCHI* promoter region. (**H-J**) Analysis of *PUCHI* expression profile by *in situ* hybridization. WT or mutant backgrounds are indicated below pictures. (Scale bars = 50 µm). Black asterisks indicate the bracts that subtend the primordia, the black arrow indicates *PUCHI* expression in the adaxial side of the boundary region between the IM and the growing primordia, and the white arrows indicate *PUCHI* signal in the abaxial side of the growing primordia. Schemes below the pictures depict the floral meristem associated with bracts in WT and *lsh1 lsh3 lsh4* mutant background. In WT background (H) the bract is cryptic and does not develop.

### Gene regulatory network establishing floral meristem boundary

Since mutations in the master floral regulator *LEAFY* (*LFY*) lead to abnormal flowers subtended by bracts (30), we wondered whether LFY could act upstream of *LSH* genes. Examination of published LFY ChIP-seq data revealed intense LFY binding to *LSH1, LSH2* and *LSH3* promoter regions (SI Appendix, Fig. S6) (31, 32). We thus examined *LSH* genes expression patterns in the *lfy-2* mutant (Fig. 3G-L) (33)and found alterations of *LSH1, LSH3* and *LSH4* expression. Specifically, *LSH1* expression became almost undetectable in the *lfy-2* mutant (compare Fig. 3A, B with G, H), whereas *LSH3* (compare Fig. 3C, D with I, J) and *LSH4* (compare Fig. 3E, F with K, L) expressions became more diffuse, ectopically present in regions such as the FM and the IM, where they are normally absent. Therefore, LFY activity is required for the localized expression of *LSH* genes in the meristem boundary and cryptic bract region, including for *LSH4* that was not directly bound by LFY in available ChIP-seq data.

In addition to *LFY*, multiple other Arabidopsis genes contribute to bract repression (Chandler, 2012) including *UNUSUAL FLORAL ORGANS* (*UFO*), *BLADE-ON-PETIOLE1 & 2* (*BOP*), *PUCHI, JAGGED* (*JAG*) and *JAGGED-like* (*JGL*)(34–38). Since BOP proteins interact with ALOG proteins in tomato and pea (11, 24), we wondered whether such interaction also occurs between Arabidopsis orthologous proteins that contribute to bract suppression. We systematically analyzed interactions between LSH proteins and several above-mentioned bract suppressors using Y2H assays (SI Appendix, Table S1). Among all proteins tested, we found that LSH1, LSH3 and LSH4 interacted with the BOP1 and BOP2 proteins (SI Appendix, Fig. S7), suggesting a conservation of the BOP-ALOG interaction in Arabidopsis.

We then exploited the LSH ampDAP-seq genomic binding profile to investigate whether some of the genes repressing bracts could be bound (and thus possibly regulated) by LSH proteins. Among the genes mentioned above, consistent LSH binding was found in the *PUCHI* promoter region (Fig. 4G and SI Appendix, Fig. S8). We thus assayed the expression of *PUCHI* by *in situ* hybridization in the *lsh1 lsh3 lsh4* triple mutant. We found that in contrast to its specific expression on the adaxial side of the WT flower meristem (Fig. 4H) (36), *PUCHI* expression was localized at the abaxial side of the early stage 1 primordia in the *lsh1 lsh3 lsh4* mutant, specifically where the bract is emerging (stage according to (39); Fig. 4I). In triple mutant floral primordia from stage 1 to stage 2, before the specification of the floral organs but when the bract has already developed, the *PUCHI* signal is mainly localized at the abaxial side of the FM (Fig. 4J). This result confirms that LSH acts upstream of *PUCHI* and suggests that the altered *PUCHI* expression could contribute to the triple *lsh1 lsh3 lsh4* mutant bract phenotype. We also checked the expression of *LFY* and *BOP1/2* genes but none of them showed an altered expression in the *lsh1 lsh3 lsh4* triple mutant (SI Appendix, Fig. S9).

## Discussion

Since the identification of the *LSH1* gene in Arabidopsis (Zhao 2004) and of the *G1* gene in rice (7) that defined the ALOG family, there has been a growing body of evidence that *ALOG* genes play important roles in many land plants (including Marchantia, Arabidopsis, tomato, rice, barley, pea or Medicago). However, fundamental questions regarding the DNA sequence they recognize, their DNA binding mechanism and their role in Arabidopsis during reproductive development have remained unanswered. In this work, we resolved several of these key questions.

First, using ampDAP-seq, we identified the DNA motif bound by ALOG proteins from Marchantia to flowering plants (Arabidopsis, tomato and rice). This result confirmed ALOG as sequence-specific DNA-binding proteins, with a bound motif conserved over a large evolutionary scale. We combined this knowledge with a set of genetic experiments showing that the Arabidopsis *lsh1 lsh3 lsh4* triple mutant flowers display a well-developed bract that is normally absent in WT flowers. Analyzing other genes involved in bract repression, we unraveled an intricate gene regulatory network downstream of the master floral regulator LFY which directly controls the proper expression pattern of both *BOP* (40–42) and *LSH* genes. As also reported in tomato and pea (11, 24), we show that Arabidopsis LSHs interact with BOP proteins. Thus, it is possible that bracts developed due to the absence of a functional LSH-BOP protein complex in the *lsh1 lsh3 lsh4* mutant. Such an LSH-BOP complex might control the expression of the bract repressor PUCHI (36). Indeed, *PUCHI* expression is no longer confined to the adaxial side of the boundary region in the *lsh1 lsh3 lsh4* triple mutant as was also observed in the *bop1 bop2* double mutant (36). The regulation of *PUCHI* by LSH is likely to be direct since we observed direct binding of the *PUCHI* promoter by LSH proteins in ampDAP-seq. These observations suggest that LSH-BOP complexes are required for the boundary identity, by confining the expression of *PUCHI* at the adaxial boundary of FM and thereby preventing bract development (Fig. 4 H-J) (36). In the *lsh1 lsh3 lsh4* triple mutant, developing bract-like organs showed chimeric features like carpelloid cells. It is thus possible that LSHs are also involved in repressing floral organ identity genes. Further genetic and molecular analyses will be needed to fully explain the mutant phenotype.

ALOG proteins have important roles in developmental transitions and reproductive development in many plants in addition to Arabidopsis. Because the ALOG motif is conserved, our work on ALOG DBD architecture and DNA binding mode is likely valid for most plant species. Whereas the organization of the helices bundle and the approximate position of the DNA-contacting surface had been roughly anticipated based on the comparison with recombinases (3), the experimental determination of the ALOG-DBD/DNA structure precisely identified residues in contact with DNA. Consistent with predictions, it showed that helices 1 and 3 but also the NLS make most DNA contacts (with an unanticipated major role of helix 3 in base recognition). The zinc ribbon does not provide additional contacts with DNA as previously thought but likely helps stabilizing the DNA-contacting helices. The ALOG-DBD/DNA structure does not resemble any classified TF/DNA structure (27, 28), and thus defined a new family in the TFClass reference classification.

In addition to the knowledge gained here, some ALOG properties remain to be investigated. For example, it was known that ALOG proteins are not interchangeable: the ALOG proteins G1 and TAW1 from rice cannot complement a Marchantia *los1* mutant while LOS1 complements a rice *g1* mutant (4). Thus, despite having similar DNA binding preferences, other features contribute to ALOG proteins’ function *in vivo*. A first explanation could be that each ALOG protein interacts with specific partners to fulfill certain functions. Another possibility is that each ALOG protein has specific biochemical properties. In fact, it was reported in tomato that ALOG proteins undergo phase separation (19, 21). This property depends on redox conditions that control the formation of intermolecular disulfide bonds, and also on intrinsic ALOG characteristics, notably their disordered regions. Cysteine residues from the zinc ribbon were proposed to contribute to the formation of TMF intermolecular disulfide bonds (19). However, our structural data suggest that mutating TMF cysteines rather disrupts the zinc ribbon (*i*.*e* abolishes DNA binding), and we did not observe any phase separation for LSH1 or LSH3 DBDs. Further work is thus required to understand the implication of each ALOG protein domain (notably non-conserved disordered regions) in their ability to undergo phase separation.

Overall, our work provides essential information on how ALOG TFs act at the molecular level. Once combined to *in vivo* DNA binding experiments, it will help understand the various roles of this important family of protein present from mosses to flowering plants.

## Materials and Methods

### Cloning

All genes were amplified from gDNA with a Phusion high fidelity polymerase (NEB) or a platinum SuperFi II polymerase (ThermoFisher) when the GC content was over 70%. All clonings were performed by Gibson Assembly and clones were checked by Sanger sequencing. Mutations were introduced by Gibson Assembly. Primers and plasmids used in this study are provided in Dataset S1 and Dataset S2, respectively.

### AmpDAP-seq

All coding sequences were cloned in the pTNT-5xmyc vector (43). We used the ampDAP-seq libraries described in (43). ampDAP-seq experiments were performed in triplicates following a previously-described protocol (44).

### ALOG recombinant protein production and purification from bacteria

All genes were cloned in the pETM-11 vector containing a N-terminal 6xHis tag and a TEV cleavage site (45). Plasmids were transformed into E.*coli* Rosetta2 (DE3) cells (Novagen). Bacteria were grown in LB medium at 37 °C up to an OD_600nm_ of 0.6. Cells were then shifted to 20 °C and 0.4 mM isopropyl b-D-1-thiogalactopyranoside (IPTG) was added. After a 3 h incubation at 20 °C, cells were collected by centrifugation and sonicated in Buffer 1 (25 mM Tris pH 8, 600 mM NaCl, 1 mM TCEP) supplemented with one EDTA-free Pierce Protease Inhibitor Tablets (ThermoFisher). Lysed cells were then centrifuged for 30 min at 15000 rpm. Supernatant was mixed with Ni Sepharose High Performance resin (Cytiva) previously equilibrated with Buffer 1. Resin was washed with Buffer 1 containing 35 mM imidazole and bound proteins were eluted with Buffer 1 containing 300 mM imidazole. Eluted proteins were mixed with TEV protease (0.01% w/w) and dialyzed overnight at 4 °C against Buffer 1.

The following day, elution was loaded again on Ni Sepharose High Performance resins (Cytiva) to remove tags and contaminants. Contaminant DNA was removed by passing proteins on Q Sepharose High Performance resin (Cytiva) pre-equilibrated with Buffer 1. DNA-free proteins (260/280 ratio below 0.6) were recovered in the flow-through and further purified by Size Exclusion Chromatography (SEC) with a Superdex 200 Increase 10/300 GL column (Cytiva) equilibrated with Buffer 2 (25 mM Tris pH 8, 150 mM NaCl, 1 mM TCEP).

### EMSA

Complementary oligos (listed in Dataset S1) were annealed overnight in annealing buffer (10 mM Tris pH 7.5, 150 mM NaCl and 1 mM EDTA). We used either TAMRA-labeled oligos (Macrogen) or complementary oligos with an overhanging G labeled with Cy5-dCTP. For this, 4 pmol of double-stranded DNA was labeled with 1 unit of Klenow fragment polymerase (NEB) and 8 pmol Cy5-dCTP (Cytiva) in Klenow buffer during 1 h at 37 °C. Enzymatic reaction was then stopped with a 10-min incubation at 65 °C. Uncropped gels are provided in Dataset S3.

Binding reactions were performed in 20 µL with different binding buffers as indicated in each figure legend. Buffer A (10 mM HEPES pH 7.5, 300 µg/mL BSA, 140 ng/µL fish sperm DNA (Sigma-Aldrich), 100 µM spermidine, 0.25 % CHAPS, 1.5 mM TCEP, 0.8 % glycerol) was used if unspecified. Otherwise, buffer B (25 mM Tris pH 8, 150 mM NaCl, 1 mM TCEP, 140 ng/µL fish sperm DNA, 0.8 % glycerol) was used as indicated in each figure legend with indicated additives. Proteins were added at indicated concentrations. All untagged proteins were purified by SEC. Binding reactions were incubated for 20 min on ice and then loaded on a 6 % native polyacrylamide gel. Gels were electrophoresed at 90 V for 75 min at 4 °C and revealed with an Amersham ImageQuant 800 imager (Cytiva).

Estimations of Kd were based on the quantification of binding experiments from Fig. 1D and SI Appendix, Fig. S2B and 2 other independent EMSAs for each protein. Kd were estimated with the Kaleidagraph software using a Michaelis-Menten model.

### co-IP

Myc- and FLAG-tagged versions of the different ALOG proteins were produced using the Quick Coupled Transcription/Translation System (TnT; Promega). 25 µL of TnT reactions producing indicated proteins were mixed with Buffer 1 to reach 150 µL and rotated at 4 °C during 1 h. 10 µL of pre-washed anti-myc beads were then added and incubated with the proteins for 1 h at 4 °C on a rotating wheel. Beads were then washed 4 times with Buffer 1. 1X protein Blue was then added to the beads, and beads were boiled for 5 min at 95 °C. Western Blots were then performed and revealed with HRP-conjugated anti-myc (Invitrogen; Cat# R951-25; 1:5000 dilution) and anti-FLAG (Sigma-Aldrich, Cat# A8592; 1:1000 dilution) antibodies. Uncropped gels are provided in Dataset S3.

### Protein crystallization, data collection and refinement

For crystallization, LSH3_S45-S190_ was purified as described above except that SEC was performed in Buffer 1 supplemented with 1 mM spermidine (Alfa Aesar). Protein was then concentrated to 5.3 mg/mL. Blunt-end complementary HPLC-purified oligos (oPR799-oPR800, see Dataset S1) were resuspended to 10 mM in 1X annealing buffer (25 mM Tris pH8, 150 mM NaCl) and annealed. Protein and DNA duplexes were directly mixed in a molar ratio of 1.1:1. ALOG-DNA was mixed at a 1:1 ratio with 0.1M HEPES, pH 7.5, 0.1M NaCl and 1.6M ammonium sulfate and crystallized using the hanging drop method. The protein-DNA complex yielded crystals after 7 days at 4 °C. Glycerol was added to the drop to ∼20% final concentration as cryoprotectant and the crystals were then flash frozen in N_2(l)_.

Diffraction data were collected at 100 K at the European Synchrotron Radiation Facility, Grenoble, France, on ID23-2 at a wavelength of 0.873 Å. Indexing was performed using MXCube (46) and the default optimized oscillation range and collection parameters used for data collection in helical collection mode to minimize radiation damage. Data were automatically processed by XDS within the Grenades pipeline (47). The data was integrated and scaled using the programs XDS and XSCALE (48). Phasing (Zn^2+^) and initial model building was performed using CRANK2 (49, 50). Model building was performed using Coot (51) and all refinements were carried out in Phenix (52). Data collection and refinement statistics are given in Table 1. The structure is deposited under PDB 8P5Q.

### Plant material and growth conditions

All experiments were performed in *Arabidopsis thaliana* accession Columbia-0 (Col-0). Plants were grown in a controlled environment at 20–22 °C either under long day conditions (16 h light/8 h dark) or under short day (8 h light/16 h dark) conditions for 4 weeks after germination and then transferred to long day conditions. When necessary, seeds of Arabidopsis were germinated on MS plates (Murashige & Skoog Medium, supplemented with 1 % sucrose and solidified with 0.7 % Plant Agar) with the appropriate selection. The *lfy-2* mutant is described in (33).

### CRISPR-Cas9 mutant generation

For generation of *lsh1, lsh3* and *lsh4* single knock-out mutants, 20-bp specific protospacers (see Dataset S1) were selected for each gene using CRISPR-P v2 database (53) and cloned into *BbsI* site of pEN-Chimera entry vector under the Arabidopsis *U6-26* promoter and then combined into the pDe-CAS9 destination vector containing the Cas9 (54) by single site Gateway® LR reaction.

### Scanning Electron Microscopy (SEM)

SEM samples were prepared as previously described (55) by gold coating them using a sputter coater (SEMPREP2; Nanotech) followed by observation with a FESEM SIGMA Scanning Electron Microscope (Zeiss).

### Yeast-Two-Hybrid (Y2H)

Coding sequences were cloned in the GAL4 system Gateway® vectors (pGADT7 and pGBKT7; Clontech) using primers listed in Dataset S1. The AH109 yeast strain was used, and yeast transformation was performed as previously described (56). Empty pGADT7 and pGBKT7 vectors were used as controls for the autoactivation. The protein-protein interaction assays were performed on selective yeast synthetic dropout medium lacking Leu, Trp, Ade, and His, and supplemented with indicated concentrations of 3-aminotriazole (3-AT). Plates were grown for 5 days at 20 °C before taking pictures. Previously-published REM35/REM35 and REM34/REM34 interactions were used as positive and negative controls, respectively (57).

### *In situ* hybridization

*In situ* hybridization analyses were performed as described in (57). Gene fragments were amplified using the primer pairs listed in Dataset S1 to generate RNA antisense probes. Evaluation of the expression profile in the inflorescence and flower meristems, which was previously published for *BOP1, BOP2, PUCHI* and *LFY* was used as a positive control (36, 58, 59). The antisense and sense RNA probes for *LSH1, LSH3* and *LSH4* were transcribed from pGEM®-T Easy with T7/SP6 RNA polymerase (Promega) according to the manufacturer’s instructions. WT reproductive meristems transversal sections hybridized with sense probes for *LSH1, LSH3* and *LSH4* are shown in SI Appendix, Fig. S10.

## Acknowledgments

We thank Miguel Blazquez for *Marchantia polymorpha* gDNA and advice and Michel Hernould for tomato gDNA. We thank Israr Ud Din, Andrea Finocchio, Bao Ngan Tu and Stefano Buratti for their technical support. We also thank Martin F. Yanofsky for providing the *AP1* coding sequence. Part of this work was carried out at NOLIMITS, an advanced imaging facility established by the Università degli Studi di Milano. This work was supported by the ANR-18-CE12-0014 ChromAuxi project to RD, the ANR-17-CE20-0014-01 Ubiflor and ANR-21-CE20-0024 Beflore projects to FP, a PhD Fellowship from CEA to PR, a post-doctoral fellowship from the University of Milan to FC, a PhD fellowships from the Doctorate School in Molecular and Cellular Biology, Università degli Studi di Milano to EF and VMB and a post-doctoral fellowship MUR - PRIN2020/2020RX4NWM to VMB.

## References

1. P. K. I. Wilhelmsson, C. Mühlich, K. K. Ullrich, S. A. Rensing, Comprehensive Genome-Wide Classification Reveals That Many Plant-Specific Transcription Factors Evolved in Streptophyte Algae. Genome Biol Evol 9, 3384–3397 (2017).

2. L. Zhao, et al., Overexpression of LSH1, a member of an uncharacterised gene family, causes enhanced light regulation of seedling development. Plant J 37, 694–706 (2004).

3. L. M. Iyer, L. Aravind, ALOG domains: provenance of plant homeotic and developmental regulators from the DNA-binding domain of a novel class of DIRS1-type retroposons. Biol Direct 7, 39 (2012).

4. S. Naramoto, Y. Hata, J. Kyozuka, The origin and evolution of the ALOG proteins, members of a plant-specific transcription factor family, in land plants. J Plant Res 133, 323–329 (2020).

5. S. Naramoto, et al., A conserved regulatory mechanism mediates the convergent evolution of plant shoot lateral organs. PLoS Biol 17, e3000560 (2019).

6. C. A. MacAlister, et al., Synchronization of the flowering transition by the tomato TERMINATING FLOWER gene. Nat Genet 44, 1393–1398 (2012).

7. A. Yoshida, T. Suzaki, W. Tanaka, H. Y. Hirano, The homeotic gene long sterile lemma (G1) specifies sterile lemma identity in the rice spikelet. Proc Natl Acad Sci U S A 106, 20103–20108 (2009).

8. X. Li, et al., TH1, a DUF640 domain-like gene controls lemma and palea development in rice. Plant Mol Biol 78, 351–359 (2012).

9. A. Yoshida, et al., TAWAWA1, a regulator of rice inflorescence architecture, functions through the suppression of meristem phase transition. Proc Natl Acad Sci U S A 110, 767–772 (2013).

10. V. M. Beretta, et al., The ALOG family members OsG1L1 and OsG1L2 regulate inflorescence branching in rice. The Plant Journal (2023) https://doi.org/10.1111/TPJ.16229 (June14, 2023).

11. L. He, et al., SYMMETRIC PETALS 1 Encodes an ALOG Domain Protein that Controls Floral Organ Internal Asymmetry in Pea (Pisum sativum L.). Int J Mol Sci 21, 4060 (2020).

12. K. Schiessl, et al., Light sensitive short hypocotyl (LSH) confer symbiotic nodule identity in the legume Medicago truncatula. bioRxiv, 2023.02.12.528179 (2023).

13. J. Zou, et al., Arabidopsis lsh8 positively regulates aba signaling by changing the expression pattern of aba-responsive proteins. Int J Mol Sci 22, 10314 (2021).

14. M. O. Press, C. Queitsch, Variability in a Short Tandem Repeat Mediates Complex Epistatic Interactions in Arabidopsis thaliana. Genetics 205, 455–464 (2017).

15. M. S. Vo Phan, I. Keren, P. T. Tran, M. Lapidot, V. Citovsky, Arabidopsis LSH10 transcription factor and OTLD1 histone deubiquitinase interact and transcriptionally regulate the same target genes. Communications Biology 2023 6:1 6, 1–11 (2023).

16. M. Lee, et al., Molecular characterization of Arabidopsis thaliana LSH1 and LSH2 genes. Genes Genomics 42, 1151–1162 (2020).

17. S. Takeda, et al., CUP-SHAPED COTYLEDON1 transcription factor activates the expression of LSH4 and LSH3, two members of the ALOG gene family, in shoot organ boundary cells. The Plant Journal 66, 1066–1077 (2011).

18. E. Cho, P. C. Zambryski, ORGAN BOUNDARY1 defines a gene expressed at the junction between the shoot apical meristem and lateral organs. Proc Natl Acad Sci U S A 108, 2154–2159 (2011).

19. X. Huang, et al., ROS regulated reversible protein phase separation synchronizes plant flowering. Nat Chem Biol 17, 549–557 (2021).

20. P. Peng, et al., The rice TRIANGULAR HULL1 protein acts as a transcriptional repressor in regulating lateral development of spikelet. Sci Rep 7, 13712 (2017).

21. X. Huang, et al., Heterotypic transcriptional condensates formed by prion-like paralogous proteins canalize flowering transition in tomato. Genome Biology 2022 23:1 23, 1–21 (2022).

22. R. C. O’Malley, et al., Cistrome and Epicistrome Features Shape the Regulatory DNA Landscape. Cell 165, 1280–1292 (2016).

23. S. J. Maerkl, S. R. Quake, A systems approach to measuring the binding energy landscapes of transcription factors. Science 315, 233–237 (2007).

24. C. Xu, S. J. Park, J. Van Eck, Z. B. Lippman, Control of inflorescence architecture in tomato by BTB/POZ transcriptional regulators. Genes Dev 30, 2048–2061 (2016).

25. L. Holm, A. Laiho, P. Törönen, M. Salgado, DALI shines a light on remote homologs: One hundred discoveries. Protein Science 32, e4519 (2023).

26. S. S. Krishna, I. Majumdar, N. V. Grishin, Structural classification of zinc fingersSURVEY AND SUMMARY. Nucleic Acids Res 31, 532–550 (2003).

27. R. Blanc-Mathieu, R. Dumas, L. Turchi, J. Lucas, F. Parcy, Plant-TFClass: a structural classification for plant transcription factors. bioRxiv, 2022.11.22.517060 (2022).

28. E. Wingender, T. Schoeps, M. Haubrock, J. Dönitz, TFClass: a classification of human transcription factors and their rodent orthologs. Nucleic Acids Res 43, D97–D102 (2015).

29. S. Bencivenga, A. Serrano-Mislata, M. Bush, S. Fox, R. Sablowski, Control of Oriented Tissue Growth through Repression of Organ Boundary Genes Promotes Stem Morphogenesis. Dev Cell 39, 198–208 (2016).

30. D. Weigel, J. Alvarez, D. R. Smyth, M. F. Yanofsky, E. M. Meyerowitz, LEAFY controls floral meristem identity in Arabidopsis. Cell 69, 843–859 (1992).

31. K. Goslin, et al., Transcription Factor Interplay between LEAFY and APETALA1/CAULIFLOWER during Floral Initiation. Plant Physiol 174, 1097–1109 (2017).

32. C. Sayou, et al., A SAM oligomerization domain shapes the genomic binding landscape of the LEAFY transcription factor. Nat Commun 7, 11222 (2016).

33. V. Grandi, V. Gregis, M. M. Kater, Uncovering genetic and molecular interactions among floral meristem identity genes in Arabidopsis thaliana. The Plant Journal 69, 881–893 (2012).

34. J. R. Dinneny, R. Yadegari, R. L. Fischer, M. F. Yanofsky, D. Weigel, The role of JAGGED in shaping lateral organs. Development 131, 1101–1110 (2004).

35. M. D. Wilkinson, G. W. Haughn, UNUSUAL FLORAL ORGANS Controls Meristem Identity and Organ Primordia Fate in Arabidopsis. Plant Cell 7, 1485–1499 (1995).

36. M. R. Karim, A. Hirota, D. Kwiatkowska, M. Tasaka, M. Aida, A Role for Arabidopsis PUCHI in Floral Meristem Identity and Bract Suppression. Plant Cell 21, 1360 (2009).

37. M. Norberg, M. Holmlund, O. Nilsson, The BLADE ON PETIOLE genes act redundantly to control the growth and development of lateral organs. Development 132, 2203–2213 (2005).

38. C. K. Ohno, G. V. Reddy, M. G. B. Heisler, E. M. Meyerowitz, The Arabidopsis JAGGED gene encodes a zinc finger protein that promotes leaf tissue development. Development 131, 1111–1122 (2004).

39. D. R. Smyth, J. L. Bowman, E. M. Meyerowitz, Early flower development in Arabidopsis. Plant Cell 2, 755–767 (1990).

40. H. Chahtane, et al., LEAFY activity is post-transcriptionally regulated by BLADE ON PETIOLE2 and CULLIN3 in Arabidopsis. New Phytologist 220, 579–592 (2018).

41. E. Moyroud, et al., Prediction of Regulatory Interactions from Genome Sequences Using a Biophysical Model for the Arabidopsis LEAFY Transcription Factor. Plant Cell 23, 1293–1306 (2011).

42. C. M. Winter, et al., LEAFY Target Genes Reveal Floral Regulatory Logic, cis Motifs, and a Link to Biotic Stimulus Response. Dev Cell 20, 430–443 (2011).

43. X. Lai, et al., The LEAFY floral regulator displays pioneer transcription factor properties. Mol Plant 14, 829–837 (2021).

44. A. Bartlett, et al., Mapping genome-wide transcription-factor binding sites using DAP-seq. Nat Protoc 12, 1659–1672 (2017).

45. A. Dümmler, A. M. Lawrence, A. de Marco, Simplified screening for the detection of soluble fusion constructs expressed in E. coli using a modular set of vectors. Microb Cell Fact 4, 34 (2005).

46. J. Gabadinho, et al., MxCuBE: a synchrotron beamline control environment customized for macromolecular crystallography experiments. J Synchrotron Radiat 17, 700–707 (2010).

47. S. Monaco, et al., Automatic processing of macromolecular crystallography X-ray diffraction data at the ESRF. J Appl Crystallogr 46, 804–810 (2013).

48. W. Kabsch, XDS. Acta Crystallogr D Biol Crystallogr 66, 125–132 (2010).

49. N. S. Pannu, et al., Recent advances in the CRANK software suite for experimental phasing. urn:issn:0907-4449 67, 331–337 (2011).

50. P. Skubak, et al., A new MR-SAD algorithm for the automatic building of protein models from low-resolution X-ray data and a poor starting model. urn:issn:2052-2525 5, 166–171 (2018).

51. P. Emsley, B. Lohkamp, W. G. Scott, K. Cowtan, Features and development of Coot. Acta Crystallogr D Biol Crystallogr 66, 486–501 (2010).

52. P. D. Adams, et al., PHENIX: a comprehensive Python-based system for macromolecular structure solution. Acta Crystallogr D Biol Crystallogr 66, 213–221 (2010).

53. H. Liu, et al., CRISPR-P 2.0: An Improved CRISPR-Cas9 Tool for Genome Editing in Plants. Mol Plant 10, 530–532 (2017).

54. F. Fauser, S. Schiml, H. Puchta, Both CRISPR/Cas-based nucleases and nickases can be used efficiently for genome engineering in Arabidopsis thaliana. Plant J 79, 348–359 (2014).

55. C. Mizzotti, M. Fambrini, E. Caporali, S. Masiero, C. Pugliesi, A CYCLOIDEA-like gene mutation in sunflower determines an unusual floret type able to produce filled achenes at the periphery of the pseudanthium. https://doi.org/10.1139/cjb-2014-0210 93, 171–181 (2015).

56. S. de Folter, R. G. H. Immink, Yeast Protein–Protein Interaction Assays and Screens. Methods in Molecular Biology 754, 145–165 (2011).

57. F. Caselli, et al., REM34 and REM35 Control Female and Male Gametophyte Development in Arabidopsis thaliana. Front Plant Sci 10, 460820 (2019).

58. S. R. Hepworth, Y. Zhang, S. McKim, X. Li, G. W. Haughn, BLADE-ON-PETIOLE–Dependent Signaling Controls Leaf and Floral Patterning in Arabidopsis. Plant Cell 17, 1434–1448 (2005).

59. S. Torti, et al., Analysis of the Arabidopsis Shoot Meristem Transcriptome during Floral Transition Identifies Distinct Regulatory Patterns and a Leucine-Rich Repeat Protein That Promotes Flowering. Plant Cell 24, 444–462 (2012).

