## Supplemental material for "The ALOG domain defines a new family of plant-specific Transcription Factors acting during Arabidopsis flower development"

**This PDF file includes:**

Figures S1 to S10
Table S1
Legends for Datasets S1 to S3
SI References

**Other supporting materials for this manuscript include the following:**

Datasets S1 to S3

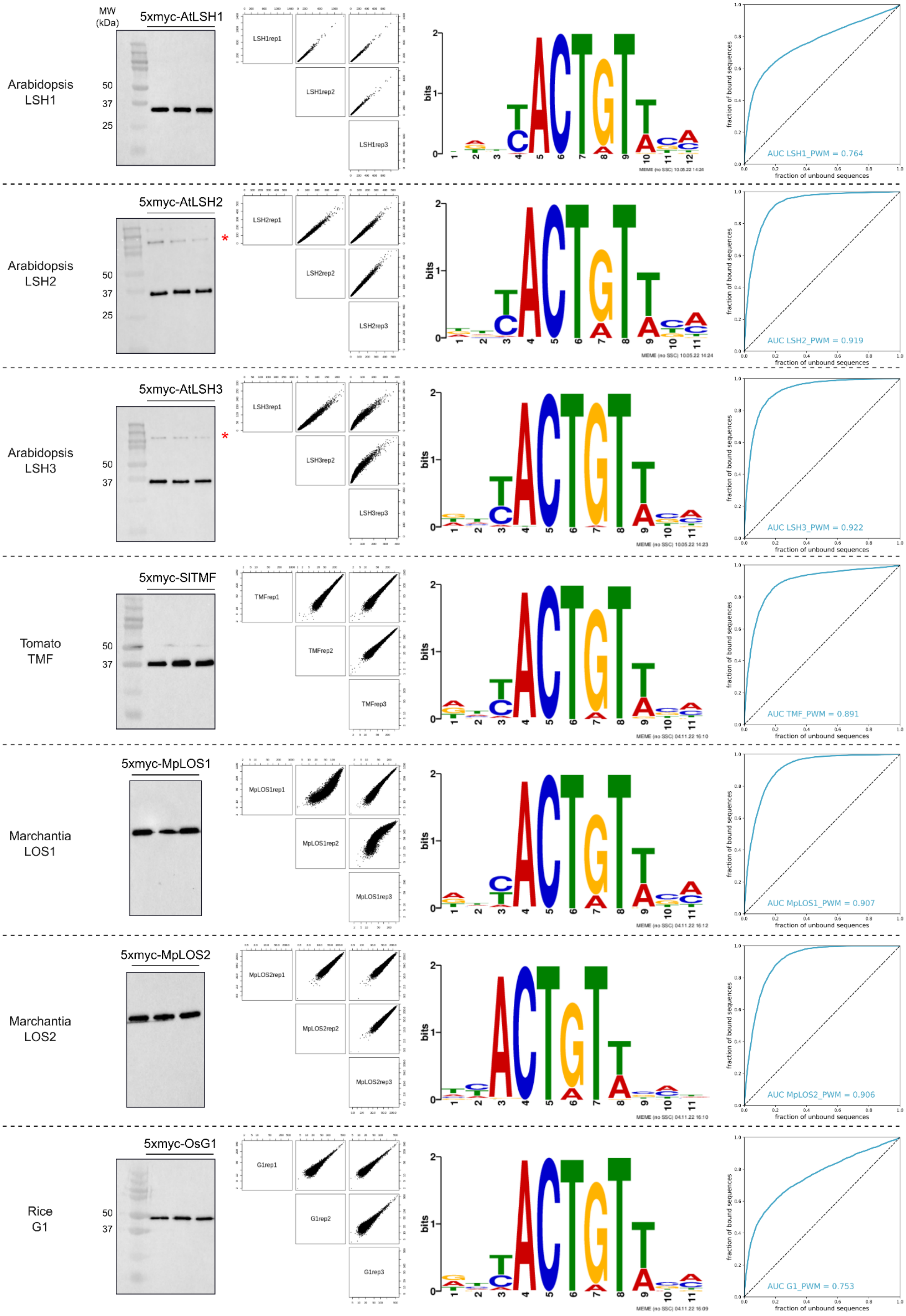

MW (kDa)
5xmyc-AtLSH1
Arabidopsis LSH1
50
37
25
LSH1rep1
LSH1rep2
LSH1rep3
bits
AUC LSH1_PWM = 0.764
fraction of bound sequences
fraction of unbound sequences
5xmyc-AtLSH2
Arabidopsis LSH2
*
LSH2rep1
LSH2rep2
LSH2rep3
AUC LSH2_PWM = 0.919
5xmyc-AtLSH3
Arabidopsis LSH3
LSH3rep1
LSH3rep2
LSH3rep3
AUC LSH3_PWM = 0.922
5xmyc-SlTMF
Tomato TMF
TMFrep1
TMFrep2
TMFrep3
AUC TMF_PWM = 0.891
5xmyc-MpLOS1
Marchantia LOS1
MpLOS1rep1
MpLOS1rep2
MpLOS1rep3
AUC MpLOS1_PWM = 0.907
5xmyc-MpLOS2
Marchantia LOS2
MpLOS2rep1
MpLOS2rep2
MpLOS2rep3
AUC MpLOS2_PWM = 0.906
5xmyc-OsG1
Rice G1
G1rep1
G1rep2
G1rep3
AUC G1_PWM = 0.753

**Fig. S1. AmpDAP-seq results for ALOG proteins from various plants.** Each line summarizes the data for one protein (see Fig. 1A). The first column shows Western Blots (WB) performed after DNA elution during ampDAP-seq experiment and revealed with an anti-myc antibody (to confirm protein binding until the last steps of the experiment for each replicate). Each lane represents one replicate. Red stars indicate probable contaminants. Molecular Weights (MW) of the protein standards are indicated (BioRad Precision Plus). Uncropped gels are provided in Dataset S3. The second column shows the experimental reproducibility of ampDAP-seq experiments through the comparison of replicate datasets 2 by 2. Each graph compares peak coverage in two different replicates. The motif generated using the 600 peaks with the strongest ampDAP-seq signal is reported in the third column. The last column represents the ROC curve using all peaks except those used to build the logo, and AUC is indicated.

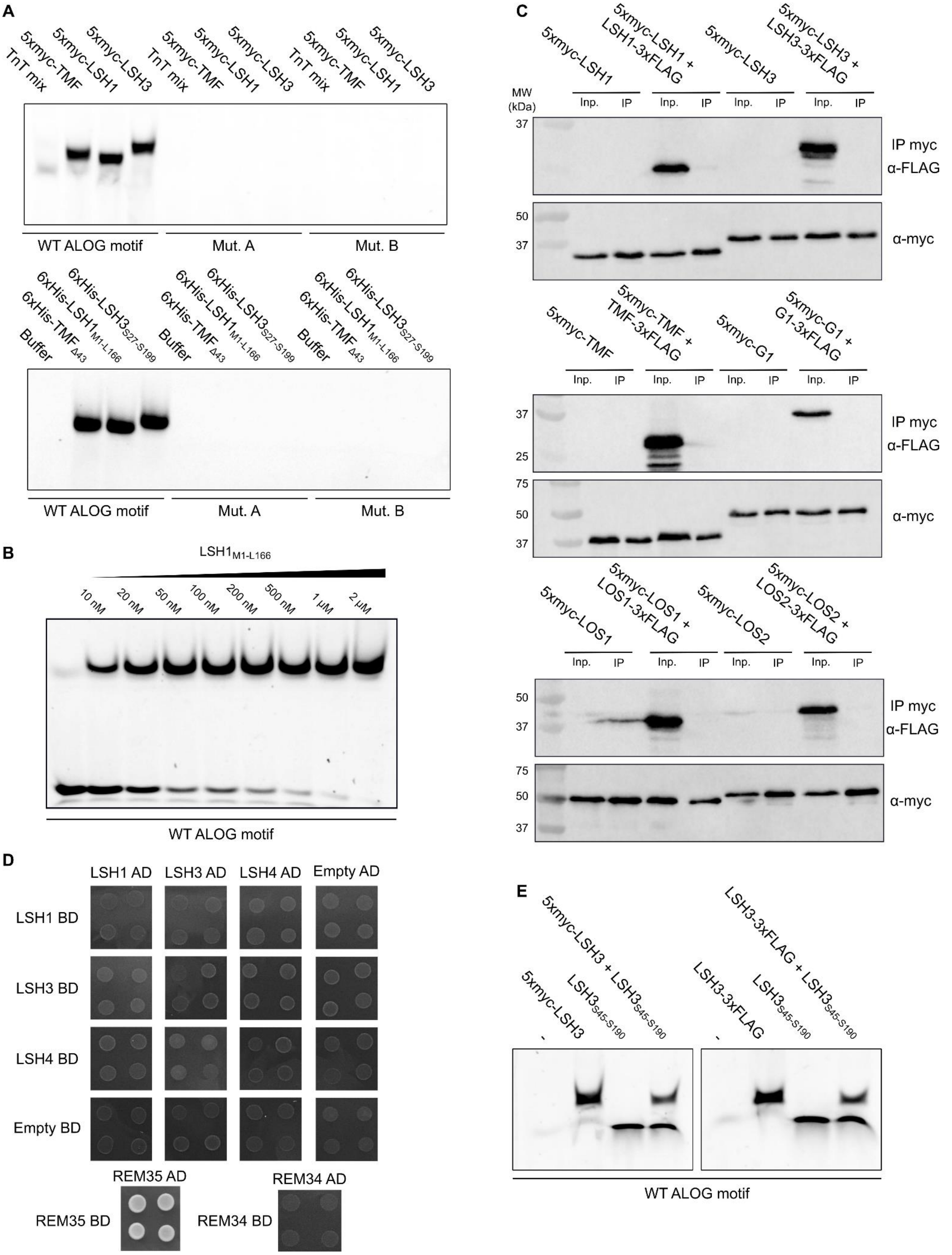

A
TnT mix
5xmyc-TMF
5xmyc-LSH1
5xmyc-LSH3
WT ALOG motif
Mut. A
Mut. B
Buffer
6xHis-TMF$_{\Delta 43}$
6xHis-LSH1$_{M1-L166}$
6xHis-LSH3$_{S27-S199}$
B
LSH1$_{M1-L166}$
10 nM
20 nM
50 nM
100 nM
200 nM
500 nM
1 µM
2 µM
WT ALOG motif
C
MW (kDa)
5xmyc-LSH1
5xmyc-LSH1 + LSH1-3xFLAG
5xmyc-LSH3
5xmyc-LSH3 + LSH3-3xFLAG
Inp.
IP
IP myc α-FLAG
α-myc
5xmyc-TMF
5xmyc-TMF + TMF-3xFLAG
5xmyc-G1
5xmyc-G1 + G1-3xFLAG
5xmyc-LOS1
5xmyc-LOS1 + LOS1-3xFLAG
5xmyc-LOS2
5xmyc-LOS2 + LOS2-3xFLAG
37
50
25
75
D
LSH1 AD
LSH3 AD
LSH4 AD
Empty AD
LSH1 BD
LSH3 BD
LSH4 BD
Empty BD
REM35 AD
REM34 AD
REM35 BD
REM34 BD
E
-
5xmyc-LSH3
LSH3$_{S45-S190}$
5xmyc-LSH3 + LSH3$_{S45-S190}$
LSH3-3xFLAG
LSH3-3xFLAG + LSH3$_{S45-S190}$
WT ALOG motif

**Fig. S2. Characterization of ALOG DBD properties.** (**A**) EMSA with indicated proteins and DNA probes. The different probes are described in Fig. 1E. 5xmyc-tagged proteins are FL and were *in vitro*-produced while 6xHis-tagged proteins are recombinant and comprise only the ALOG domain. $TMF_{\Delta 43}$ corresponds to a TMF truncated version lacking the first 43 aa. (**B**) EMSA with ALOG highest-score sequence DNA probe and $LSH1_{M1\text{-}L166}$. Based on the analysis of 3 independent EMSAs, we found an apparent Kd of 29 nM for LSH1-DBD/DNA. (**C**) co-IP with indicated *in vitro*-produced proteins. Molecular Weights (MW) of the protein standards are indicated (BioRad Precision Plus). (**D**) Interaction between LSH1-4 tested by Y2H (–L-W-H + 2.5 mM 3-AT selective media). Empty pGADT7 and pGBKT7 vectors were used as controls for the autoactivation. Previously published REM35/REM35 and REM34/REM34 interactions were used as positive and negative controls, respectively (1). (**E**) EMSA with indicated proteins and ALOG highest-score sequence DNA probe. 5xmyc-LSH3 and LSH3-3xFLAG are *in vitro*-produced FL versions while $LSH3_{S45\text{-}S190}$ is recombinant. When two LSH3 versions were mixed, the amount of each protein was half the amount used for reactions with the protein alone. Note that no LSH3 heterodimer is observed. Uncropped gels are provided in Dataset S3.

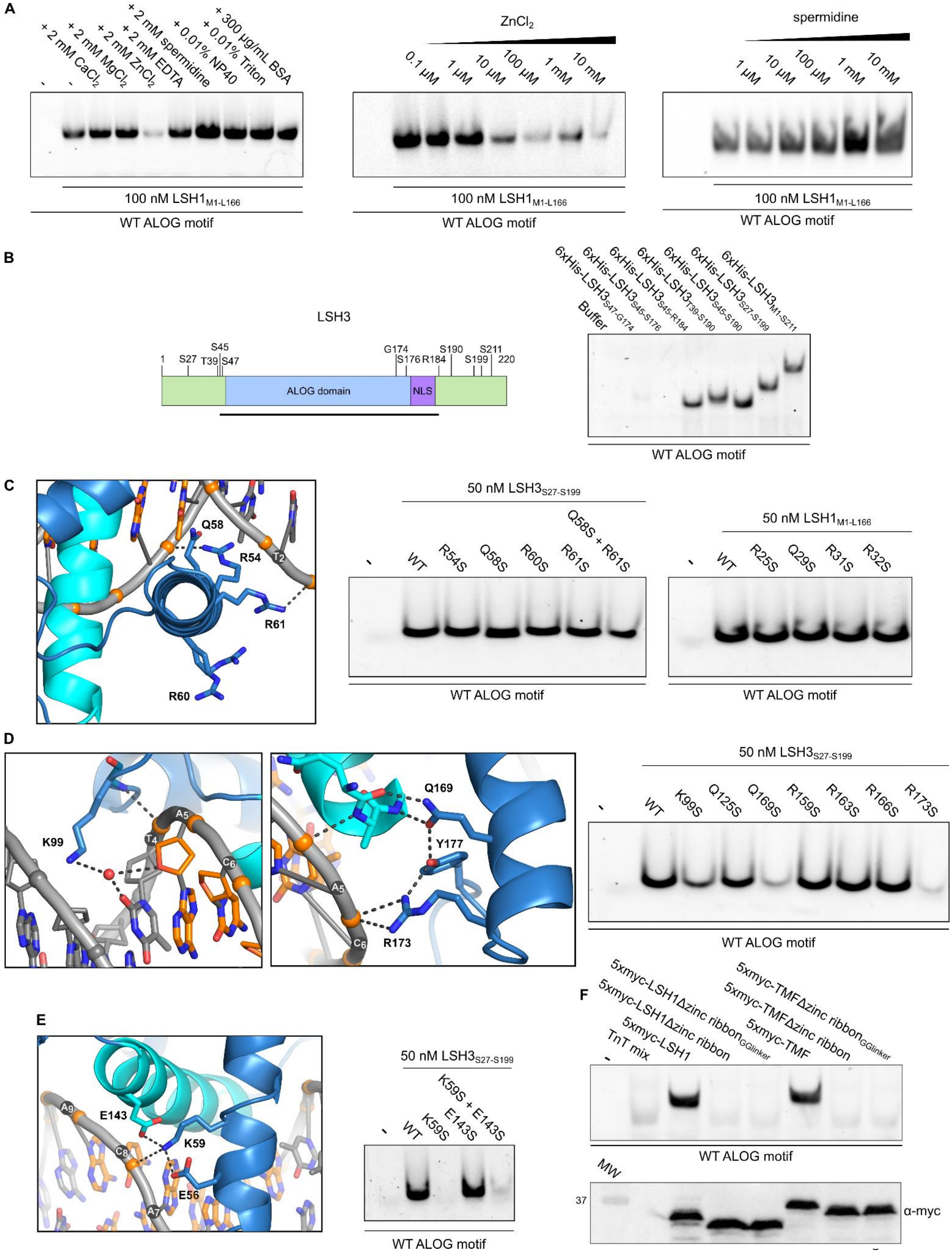

A
-
+ 2 mM $CaCl_2$
-
+ 2 mM $MgCl_2$
+ 2 mM $ZnCl_2$
+ 2 mM EDTA
+ 2 mM spermidine
+ 0.01% NP40
+ 0.01% Triton
+ 300 μg/mL BSA
100 nM LSH1$_{M1-L166}$
WT ALOG motif
$ZnCl_2$
0.1 μM
1 μM
10 μM
100 μM
1 mM
10 mM
100 nM LSH1$_{M1-L166}$
WT ALOG motif
spermidine
1 μM
10 μM
100 μM
1 mM
10 mM
100 nM LSH1$_{M1-L166}$
WT ALOG motif
B
LSH3
1
S27
T39
S45
S47
G174
S176
R184
S190
S199
S211
220
ALOG domain
NLS
Buffer
6xHis-LSH3$_{S47-G174}$
6xHis-LSH3$_{S45-S176}$
6xHis-LSH3$_{S45-R184}$
6xHis-LSH3$_{T39-S190}$
6xHis-LSH3$_{S45-S190}$
6xHis-LSH3$_{S27-S199}$
6xHis-LSH3$_{M1-S211}$
WT ALOG motif
C
Q58
R54
T2
R61
R60
50 nM LSH3$_{S27-S199}$
-
WT
R54S
Q58S
R60S
R61S
Q58S + R61S
WT ALOG motif
50 nM LSH1$_{M1-L166}$
-
WT
R25S
Q29S
R31S
R32S
WT ALOG motif
D
K99
A5
T4
C6
Q169
Y177
A5
C6
R173
50 nM LSH3$_{S27-S199}$
-
WT
K99S
Q125S
Q169S
R159S
R163S
R166S
R173S
WT ALOG motif
E
A9
E143
C8
K59
E56
A7
50 nM LSH3$_{S27-S199}$
-
WT
K59S
E143S
K59S + E143S
WT ALOG motif
F
-
TnT mix
5xmyc-LSH1
5xmyc-LSH1Δzinc ribbon
5xmyc-LSH1Δzinc ribbon$_{GGlinker}$
5xmyc-TMF
5xmyc-TMFΔzinc ribbon
5xmyc-TMFΔzinc ribbon$_{GGlinker}$
WT ALOG motif
MW
37
α-myc

**Fig. S3. Characterization of ALOG DNA binding properties.** (**A**) Effect of several additives on LSH3 DNA binding *in vitro*. Buffer B (see Methods) was supplemented with indicated compounds (left). $ZnCl_2$ (middle) and spermidine (right) were further studied with a concentration range. (**B**) Determination of the minimal LSH3 domain sufficient for DNA binding. Schematic of LSH3 (left). The black bar represents the minimal LSH3 domain for DNA binding. Drawing is not at scale. EMSA with indicated truncated versions (right). Protein concentrations = 100 nM. Each protein was mixed with the same DNA-containing mix. (**C**) LSH3 DBD helix 1 residues are not essential for DNA binding. Close-up view of LSH3 helix 1 residues (left). EMSA with indicated proteins and ALOG WT probe (middle and right). (**D**) Role of other conserved LSH3 DBD residues in DNA binding. Close-up view of K99 (left) and helix 4 residues (middle). EMSA with indicated proteins and ALOG highest-score sequence DNA probe (right). Mutations of K99, Q169 and R173 reduce the DNA binding. (**E**) Role of K59 in DNA binding. Close-up view of K59 interactions (left). EMSA with indicated proteins and ALOG WT probe (right). K59 is essential for DNA binding. (**F**) EMSA with indicated proteins and ALOG highest-score sequence DNA probe (top). LSH1 and TMF zinc ribbons were either deleted (Δzinc ribbon) or replaced by a GG linker found in the XerD recombinase (Δzinc $\text{ribbon}_{\text{GGlinker}}$). The production of the different 5xmyc-tagged proteins used in EMSA was confirmed by Western Blot (bottom). The linker does not restore DNA binding. Uncropped gels are provided in Dataset S3.

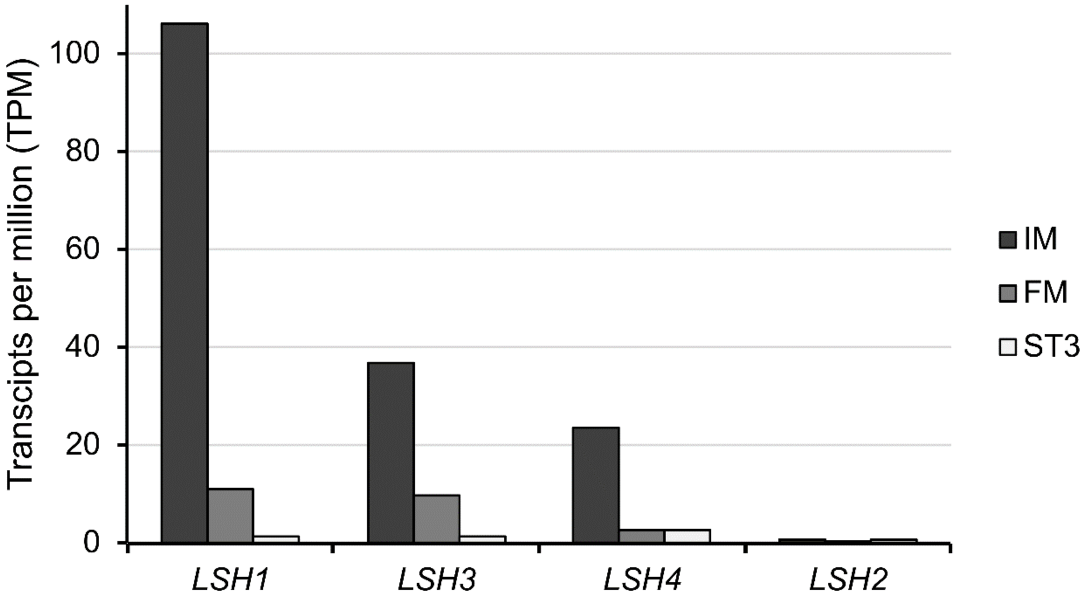


**Fig. S4. Expression of *LSH1-4* in different reproductive tissues.** RNA-seq data are from (2). Read counts are presented in transcripts per million. IM = Inflorescence Meristem, FM = Flower Meristem, ST3 = flower primordia at stage 3.

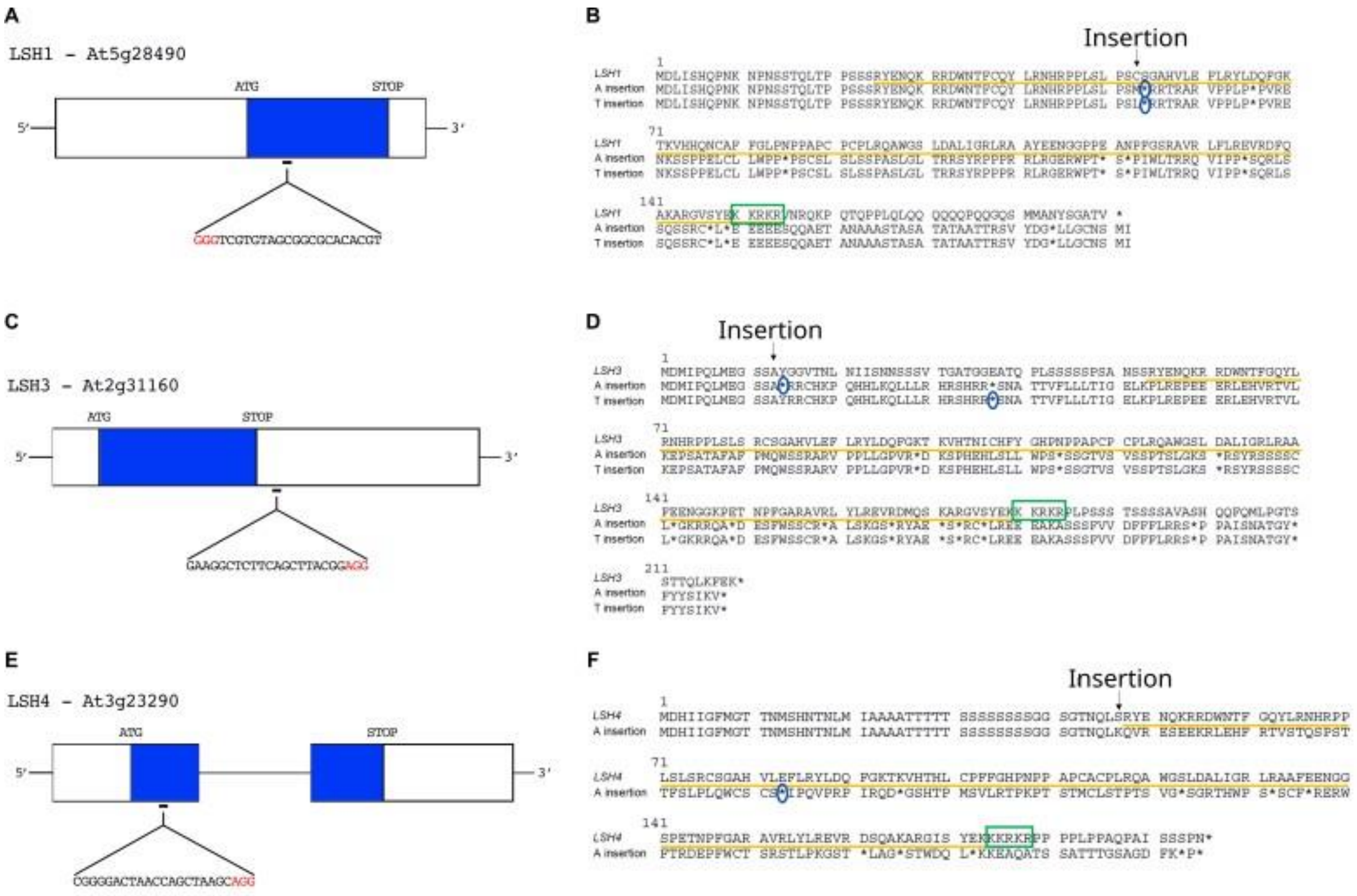


**Fig. S5. Mutation of *LSH1*, *LSH3* and *LSH4* by CRISPR-Cas9.** (**A, C, E**) *LSH1, LSH3* and *LSH4* gene models. Exons (blue rectangle), introns (black line), and 5' and 3' UTRs (white rectangle) are represented. The position and the target sequence of the sgRNA, which targets *LSH1* exon, 153 bp downstream the ATG site (A); LSH3 exon, 24 bp downstream the ATG site (C) and LSH4 first exon, 121 bp downstream the ATG site (E) are indicated. The PAM sequence is indicated in red. (**B, D, F**) Amino acid sequence alignments of WT and mutated LSH versions. The different base pair insertions in mutants are indicated on the left. The ALOG domain is underlined in yellow, the green rectangles indicate the NLS and premature stop codons (*) are circled in blue. In all mutants, insertions cause the formation of premature stop codons (ALOG domain and NLS disrupted).

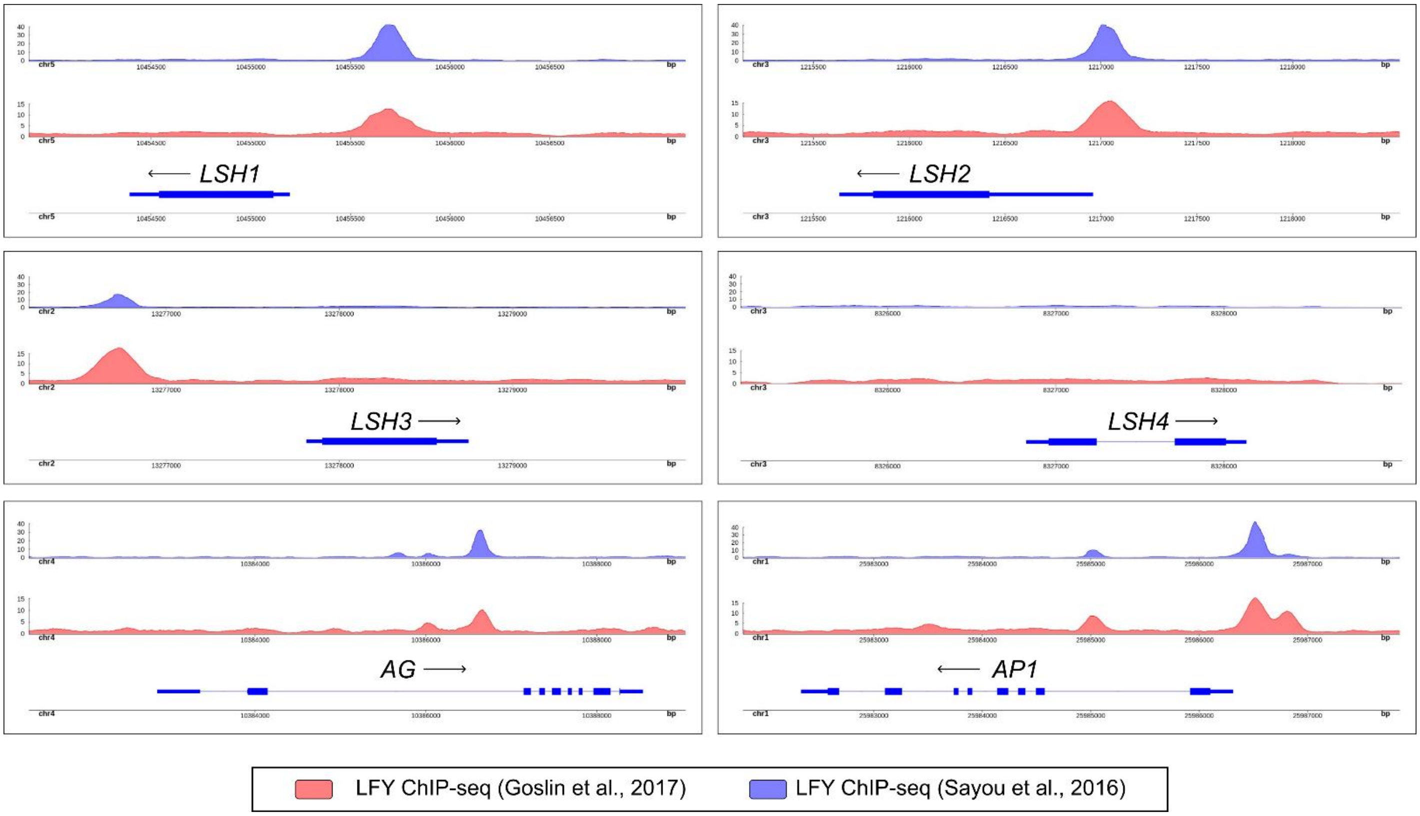


**Fig. S6. LFY binding to *LSH* regions.** LFY ChIP-seq signal from *35S::LFY-GR ap1 cal* inflorescences (3) or *35S::LFY* seedlings (4) on *LSH1-4* genes, as well as on the well-characterized LFY targets *APETALA1* and *AGAMOUS* for comparison.

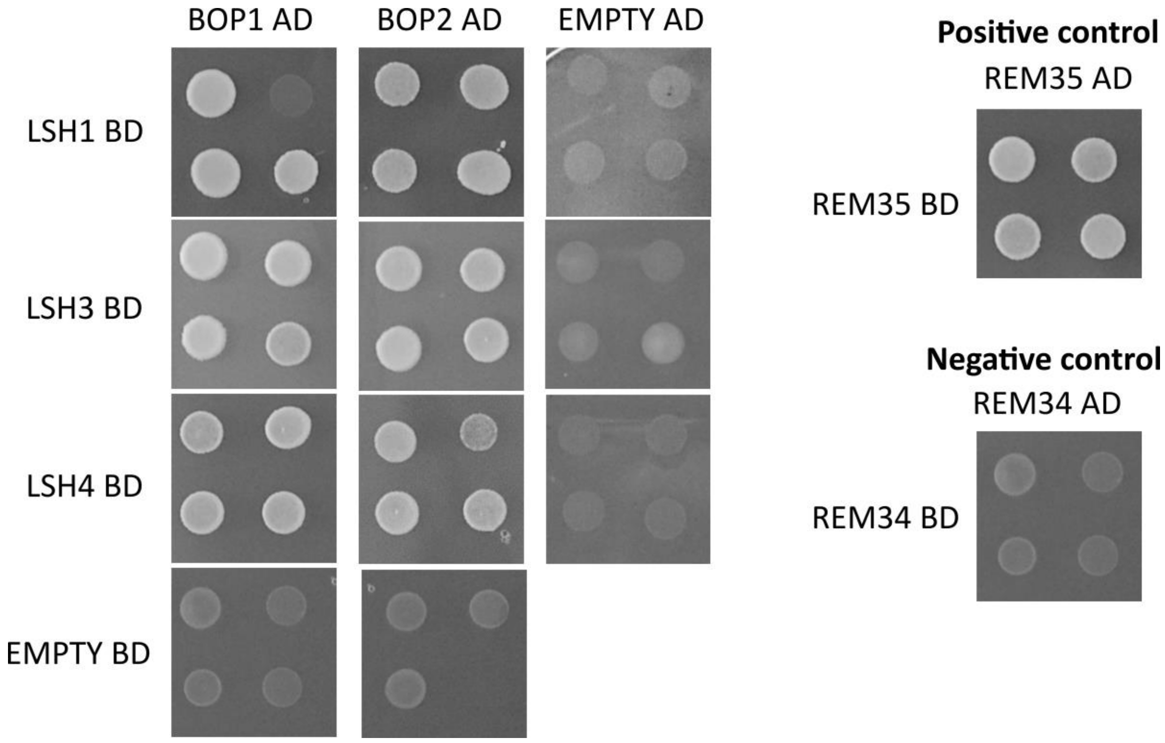


**Fig. S7. LSH1, LSH3 and LSH4 interact with BOP1 and BOP2.** Yeast-two-hybrid (Y2H) assay showing the interactions between LSH1, LSH3, LSH4 and BOP1 and BOP2, on –L-W-H + 2.5 mM 3-AT selective media. Empty pGADT7 and pGBKT7 vectors were used as controls for the autoactivation. As positive and negative controls, the already published interactions between REM35 and REM35; and between REM34 and REM34 respectively were used (1).

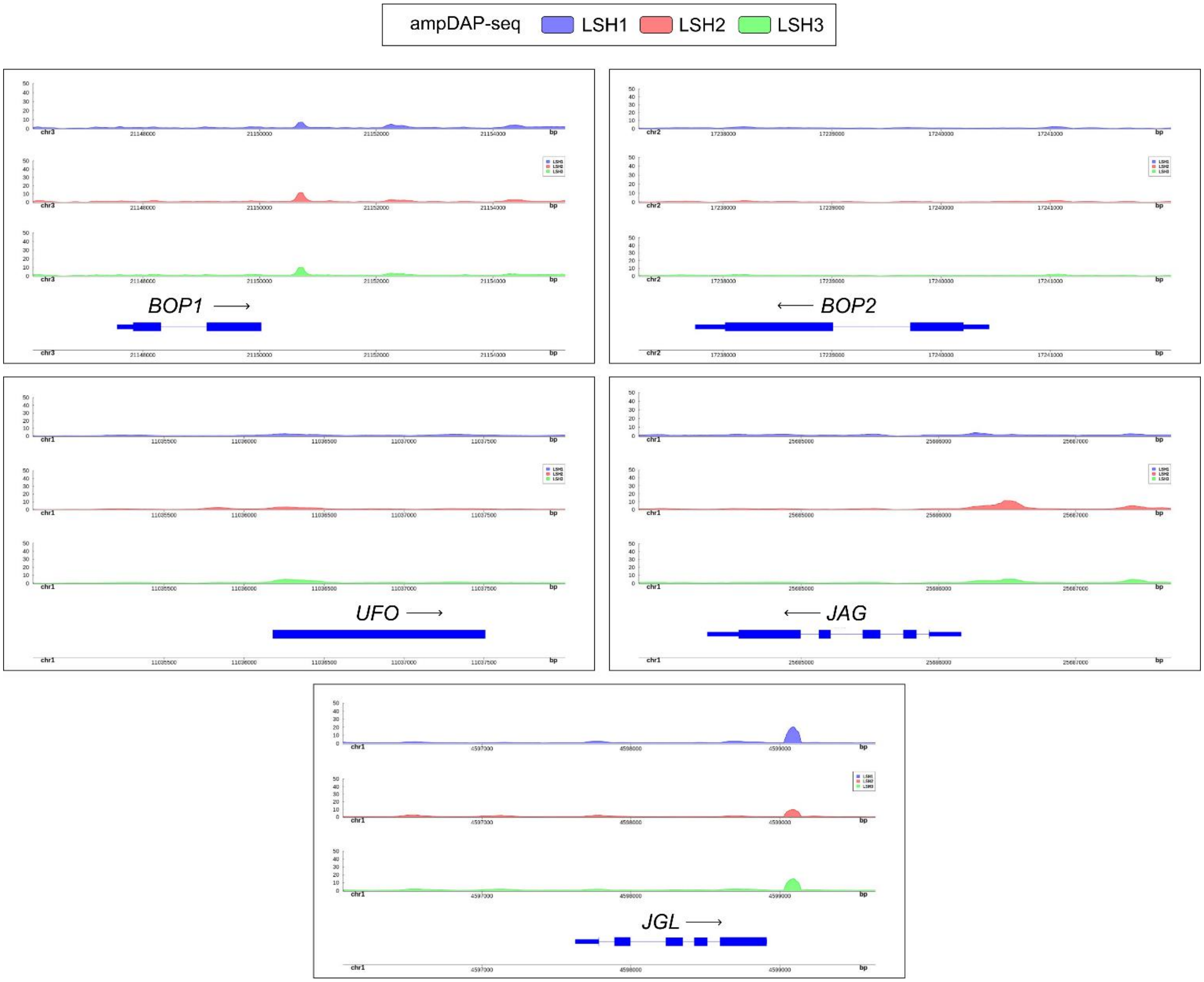


**Fig. S8. LSH binding to bract repressors genomic regions.** Genome browser view showing ampDAP-seq binding of LSH proteins to *BOP1*, *BOP2*, *UFO*, *JAG* and *JGL* genomic regions. None of these regions show clear LSH peaks.

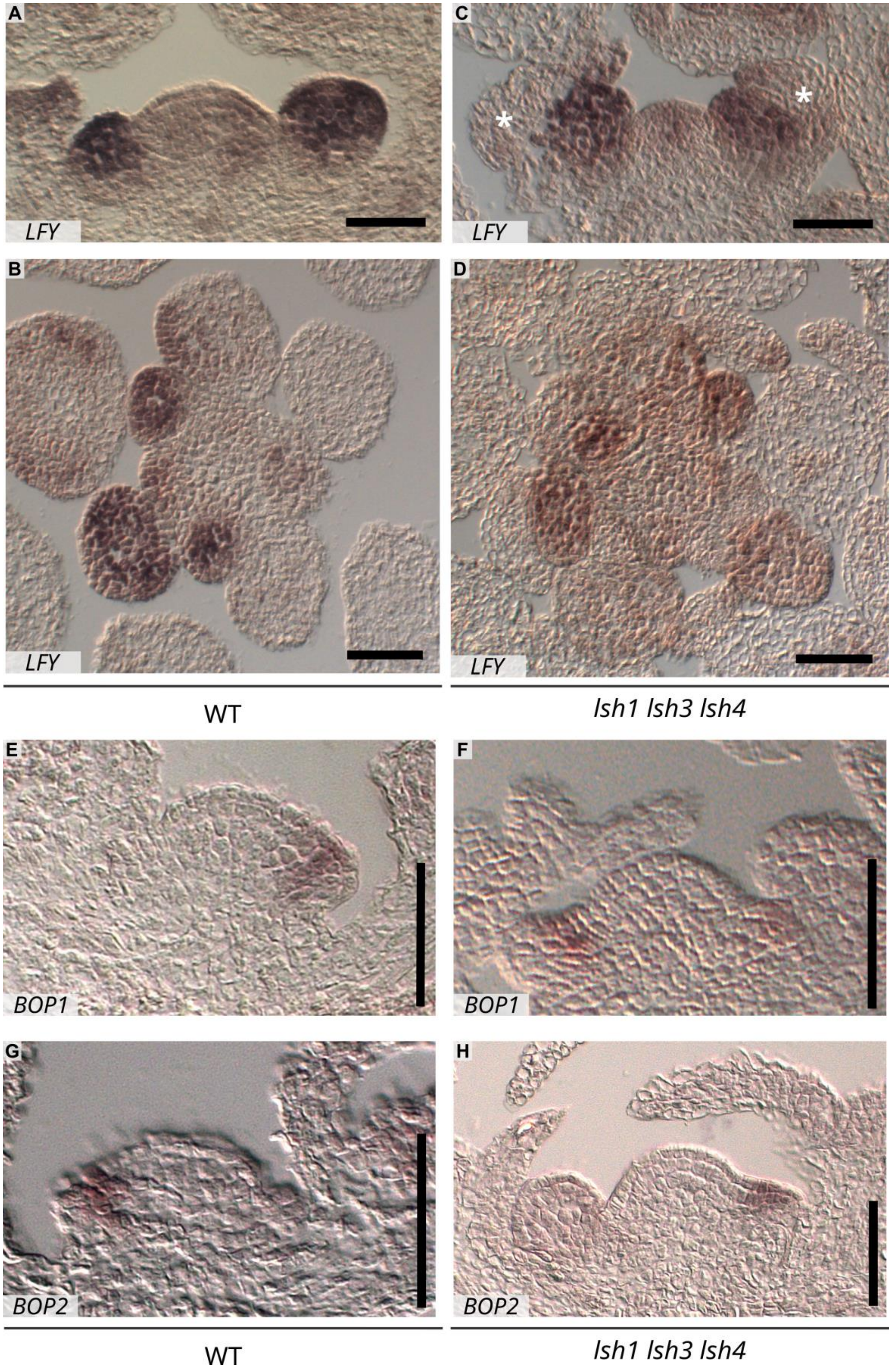
A
LFY
B
LFY
C
*
*
LFY
D
LFY
WT
lsh1 lsh3 lsh4
E
BOP1
F
BOP1
G
BOP2
H
BOP2
WT
lsh1 lsh3 lsh4

**Fig. S9. Expression of LFY and BOP1/2 RNAs in WT and *lsh1 lsh3 lsh4* backgrounds.** Expression profile of *LFY* (A-D), *BOP1* (E, F) and *BOP2* (G, H) were analyzed by *in situ* hybridization in WT (A, B, E, G) and *lsh1 lsh3 lsh4* (C, D, F, H) reproductive tissues. A, C, E, F are longitudinal sections, B, D, G, H are transversal sections. (Scale bars = 50 μm). White asterisks indicate the bracts subtending the primordia.

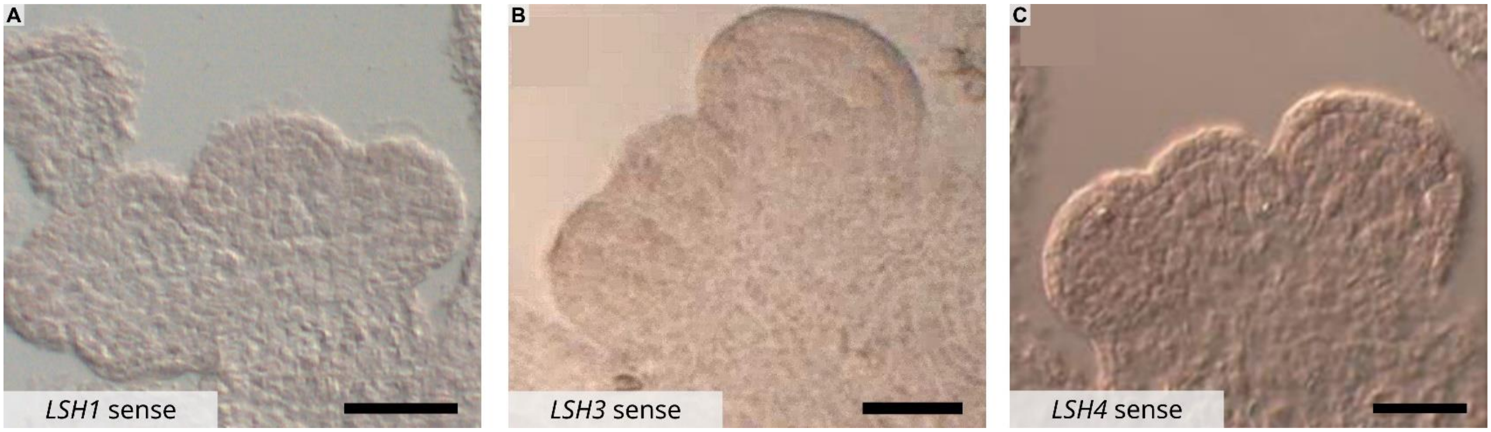


**Fig. S10. Negative controls for *in situ* hybridization experiments.** WT reproductive meristems longitudinal sections hybridized with the sense negative control probes for *LSH1* (A), *LSH3* (B) and *LSH4* (C; Scale bars = 50 μm).

| Interactions tested | -L-W | -L-W-H+ 2.5 mM 3AT | -L-W-H+ 5 mM 3AT | -L-W-H+ 10 mM3AT |
|---|---|---|---|---|
| pAD-AP1 + pBD-LSH1 | + | - | - | - |
| pAD-AP1 + pBD-LSH3 | + | - | - | - |
| pAD-AP1 + pBD-LSH4 | + | - | - | - |
| **pAD-AP1 + pBD-EMPTY** | **+** | - | - | - |
| pAD-LSH1 + pBD-LFY | + | - | - | - |
| pAD-LSH3 + pBD-LFY | + | - | - | - |
| pAD-LSH4 + pBD-LFY | + | - | - | - |
| **pAD-EMPTY + pBD-LFY** | **+** | - | - | - |
| pAD-PUCHI + pBD-LSH1 | + | - | - | - |
| pAD-PUCHI + pBD-LSH3 | + | - | - | - |
| pAD-PUCHI + pBD-LSH4 | + | - | - | - |
| pAD-LSH1 + pBD-PUCHI | + | + | + | + |
| pAD-LSH3 + pBD-PUCHI | + | + | + | + |
| pAD-LSH4 + pBD-PUCHI | + | + | + | + |
| **pAD-PUCHI + pBD-EMPTY** | **+** | - | - | - |
| **pAD-EMPTY + pBD-PUCHI** | **+** | **+** | **+** | **+** |
| pAD-BOP1 + pBD-LSH1 | + | + | + | + |
| pAD-BOP1 + pBD-LSH3 | + | + | + | + |
| pAD-BOP1 + pBD-LSH4 | + | + | + | + |
| pAD-LSH1 + pBD-BOP1 | + | + | + | + |
| pAD-LSH3 + pBD-BOP1 | + | + | + | + |
| pAD-LSH4 + pBD-BOP1 | + | + | + | + |
| **pAD-BOP1 + pBD-EMPTY** | **+** | - | - | - |
| **pAD-EMPTY + pBD-BOP1** | **+** | **+** | **+** | **+** |
| pAD-BOP2 + pBD-LSH1 | + | + | + | + |
| pAD-BOP2 + pBD-LSH3 | + | + | + | + |
| pAD-BOP2 + pBD-LSH4 | + | + | + | + |
| pAD-LSH1 + pBD-BOP2 | + | + | + | + |
| pAD-LSH3 + pBD-BOP2 | + | + | + | + |
| pAD-LSH4 + pBD-BOP2 | + | + | + | + |
| **pAD-BOP2 + pBD-EMPTY** | **+** | - | - | - |
| **pAD-EMPTY + pBD-BOP2** | **+** | **+** | **+** | **+** |
| pAD-JAG + pBD-LSH1 | + | - | - | - |
| pAD-JAG + pBD-LSH3 | + | - | - | - |
| pAD-JAG + pBD-LSH4 | + | - | - | - |
| pAD-LSH1 + pBD-JAG | + | - | - | - |
| pAD-LSH3 + pBD-JAG | + | - | - | - |
| pAD-LSH4 + pBD-JAG | + | - | - | - |
| **pAD-JAG + pBD-EMPTY** | **+** | - | - | - |
| **pAD-EMPTY + pBD-JAG** | **+** | - | - | - |
| pAD-JGL + pBD-LSH1 | + | - | - | - |
| pAD-JGL + pBD-LSH3 | + | - | - | - |
| pAD-JGL + pBD-LSH4 | + | - | - | - |
| pAD-LSH1 + pBD-JGL | + | - | - | - |
| pAD-LSH3 + pBD-JGL | + | - | - | - |
| pAD-LSH4 + pBD-JGL | + | - | - | - |
| **pAD-JGL + pBD-EMPTY** | **+** | - | - | - |
| **pAD-EMPTY + pBD-JGL** | **+** | - | - | - |
| pAD-LSH1 + pBD-LSH1 | + | - | - | - |
| pAD-LSH1 + pBD-LSH3 | + | - | - | - |
| pAD-LSH1 + pBD-LSH4 | + | - | - | - |
| pAD-LSH1 + pBD-LSH1 | + | - | - | - |
| pAD-LSH3 + pBD-LSH1 | + | - | - | - |
| pAD-LSH4 + pBD-LSH1 | + | - | - | - |
| **pAD-LSH1 + pBD-EMPTY** | **+** | - | - | - |
| **pAD-EMPTY + pBD-LSH1** | **+** | - | - | - |
| pAD-LSH3 + pBD-LSH1 | + | - | - | - |
| pAD-LSH3 + pBD-LSH3 | + | - | - | - |
| pAD-LSH3 + pBD-LSH4 | + | - | - | - |
| pAD-LSH1 + pBD-LSH3 | + | - | - | - |
| pAD-LSH3 + pBD-LSH3 | + | - | - | - |
| pAD-LSH4 + pBD-LSH3 | + | - | - | - |
| **pAD-LSH3 + pBD-EMPTY** | **+** | - | - | - |
| **pAD-EMPTY + pBD-LSH3** | **+** | - | - | - |
| pAD-LSH4 + pBD-LSH1 | + | - | - | - |
| pAD-LSH4 + pBD-LSH3 | + | - | - | - |
| pAD-LSH4 + pBD-LSH4 | + | - | - | - |
| pAD-LSH1 + pBD-LSH4 | + | - | - | - |
| pAD-LSH3 + pBD-LSH4 | + | - | - | - |
| pAD-LSH4 + pBD-LSH4 | + | - | - | - |
| **pAD-LSH4 + pBD-EMPTY** | **+** | - | - | - |
| **pAD-EMPTY + pBD-LSH4** | **+** | - | - | - |
| **pAD-REM35 + pBD-REM35** | + | + | + | + |
| **pAD-REM34 + pBD-REM34** | + | - | - | - |

**Table S1. Summary of Y2H interactions.** Yeasts were grown on -L-W media and on -L-W-H selective media supplemented with indicated concentrations of 3-AT. (+) and (-) indicate positive or negative interactions. In bold are indicated the tests with empty pGADT7 and pGBKT7 vectors as controls for the autoactivation. Previously published REM35/REM35 and REM34/REM34 interactions were used as positive and negative controls, respectively (green; (1)).

**Dataset S1 (separate file).** List of primers used in this study

**Dataset S2 (separate file).** List of plasmids used in this study

**Dataset S3 (separate file).** Uncropped gels

**SI References**


1. F. Caselli, *et al.*, REM34 and REM35 Control Female and Male Gametophyte Development in Arabidopsis thaliana. *Front Plant Sci* **10**, 460820 (2019).
2. O. Mantegazza, *et al.*, Gene coexpression patterns during early development of the native Arabidopsis reproductive meristem: novel candidate developmental regulators and patterns of functional redundancy. *The Plant Journal* **79**, 861–877 (2014).
3. K. Goslin, *et al.*, Transcription Factor Interplay between LEAFY and APETALA1/CAULIFLOWER during Floral Initiation. *Plant Physiol* **174**, 1097–1109 (2017).
4. C. Sayou, *et al.*, A SAM oligomerization domain shapes the genomic binding landscape of the LEAFY transcription factor. *Nat Commun* **7**, 11222 (2016).
